# Medial preoptic area FoxO1 controls metabolic adaptation in a sexually dimorphic manner

**DOI:** 10.1101/2025.06.25.661575

**Authors:** Pei Luo, Xiaohua Yang, Qi Xu, Lau Lee How, Valeria C. Torres Irizarry, Leslie Carrillo-Saenz, Marcos David Munoz, Wei Dong, Lucas Ibrahimi, Sarah Schaul, Nirali Patel, Nimisha Antony, Devin Dixit, Maya Kota, Christopher Freeman, Elias L Cruz, Chunmei Wang, Graziano Pinna, Yuwei Jiang, Chong Wee Liew, Gang Shu, Hui Ye, Yanlin He, Terry Unterman, Pingwen Xu

## Abstract

The medial preoptic area (MPOA) of the hypothalamus is essential for metabolic adaptation to environmental challenges, though the molecular mechanisms underlying this process remain poorly understood. Here, we investigate the role of Forkhead transcription factor O1 (FoxO1), a key mediator of stress adaptation, in MPOA-dependent metabolic responses to temperature and nutritional changes. Our findings reveal sex-specific responses to both nutritional and temperature challenges. In female mice, but not males, a high-fat diet (HFD) challenge decreased FoxO1 expression in the MPOA. Specific deletion of FoxO1 in MPOA neurons (FoxO1-KO^MPOA^) had no effect on body weight under normal chow-fed conditions but protected females from HFD-induced obesity (DIO). These protected females exhibited increased lean mass, decreased fat mass, enhanced thermogenesis, increased energy expenditure, and reduced food intake under HFD conditions. They also showed enhanced cold-induced heat production at 6°C, though this effect vanished at thermoneutrality (30°C). The protection against DIO was abolished by ovariectomy (OVX) and was not restored by 17β-estradiol supplementation, suggesting an estrogen-independent mechanism. Conversely, constitutive activation of FoxO1 in MPOA neurons (FoxO1-CA^MPOA^) increased DIO susceptibility in both sexes. Together, these findings demonstrate that FoxO1^MPOA^ plays a crucial role in coordinating metabolic adaptation to nutritional and temperature challenges specifically in female mice.

## Introduction

A major challenge in today’s modern society is the evolutionary adaptation of human metabolic control systems to conditions of food scarcity and stable environmental temperature, which conflicts with the current environment of food abundance and global warming due to technological advancements and modernization^1–6^. This mismatch has led to a rapidly increasing epidemic of obesity and other metabolic diseases. Understanding how the central nervous system manages metabolic adaptation to temperature and nutritional challenges is essential. This knowledge may help develop therapies targeting metabolic syndrome, particularly relevant in today’s society.

Many proteins help maintain whole-body metabolic balance under stress conditions, including the transcription factor forkhead proteins, especially FoxO1^7^. The FoxO1 signaling pathway was first discovered in *Caenorhabditis elegans* (*C. elegans*). Two independent research groups identified DAF-16 (a homolog of mammalian FoxO1) as a negative modulator of DAF-2 (a homolog of the mammalian insulin receptor) signaling in *C. elegans*^8,9^. By the end of the last century, it was clear that the stress resistance of the mutant DAF-2 *C. elegans* depend on the DAF-16 gene^10^. This indicates that DAF-16/ FoxO1 may play an essential role in stress response. Further studies using mouse models have confirmed FoxO1 as a key mediator of metabolic adaptation to fasting and cold exposure. During fasting, FoxO1 in the liver drives gluconeogenesis, while FoxO1 in the muscle promotes a metabolic shift from glucose catabolism to lipid oxidation to maintain glucose levels^11–14^. Additionally, FoxO1 has recently been identified as a key transcription factor for autonomous cold adaptation. Specifically, cold-induced FoxO1 nuclear transport supports cold survival and tissue storage^15^. Therefore, FoxO1 plays a vital role in metabolic adaptation to nutritional and temperature challenges, ensuring that cells, organs, and ultimately the whole organism maintain resilience and systemic homeostasis.

Interestingly, FoxO1 expressed by hypothalamic pro-opiomelanocortin (POMC) neurons^16,17^, agouti-related peptide (AgRP) neurons^18,19^, steroidogenic factor 1 (SF-1) neurons^20^, and mesolimbic dopamine neurons^21^ has been recently reported to modulate systemic metabolism by regulating food intake, energy expenditure, and leptin/insulin sensitivity. This suggests a vital role for brain FoxO1 in energy homeostasis regulation. However, it is still unknown whether brain FoxO1 contributes to metabolic adaptation to nutritional and temperature challenges.

The medial preoptic area (MPOA), an anterior part of the hypothalamus, is a temperature sensory node and control center for body temperature regulation, which modulates food intake and energy expenditure in response to ambient temperature fluctuation^22–24^. Temperature-sensitive MPOA neurons respond to ambient temperature changes to increase or decrease energy expenditure during cold or warm exposure to maintain temperature homeostasis^24,25^. Notably, these MPOA neurons also counterbalance the changed energy expenditure levels by adjusting the amount of food consumption to prevent body weight loss (cold temperature) or gain (warm temperature)^24,26,27^. Recently, MPOA neurons have also been shown to coordinate energy deficiency-induced hypothermic and hypometabolic responses known as torpor^28–30^, indicating a role of MPOA neurons in the thermoregulatory and metabolic responses to energy deficiency. Thus, the MPOA-driving interaction between energy expenditure and food intake ensures temperature and energy homeostasis during nutritional and temperature challenges. It is worth investigating whether FoxO1 is expressed in the MPOA and, if so, whether it contributes to the metabolic adaptations mediated by MPOA neurons.

In this study, we investigated the expression pattern of FoxO1 in the MPOA (FoxO1^MPOA^) and its metabolic physiological relevance. Building on previous observations that FoxO1 plays a crucial role in resistance to metabolic stresses^7^ and MPOA is vital for metabolic adaptation to temperature challenges and fasting^24,31^, we further assessed the potential role of FoxO1^MPOA^ in metabolic adaptation to nutritional and temperature stresses in mice.

## Results

### FoxO1^MPOA^ responds to the high-fat diet (HFD) challenge in a sexually dimorphic way

To investigate whether FoxO1 is involved in MPOA neuronal response to nutritional challenges, we assessed FoxO1 expression levels in the MPOA of wild-type C57BL/6J mice (8 weeks old) after 2 weeks on either a chow diet or HFD. The HFD challenge significantly decreased MPOA FoxO1 mRNA expression levels in female mice but not in male mice (Fig. 1A). Similarly, we observed decreased FoxO1-positive neurons in the MPOA of female mice, but not male mice, after 7 days of HFD feeding, though not after 3 days (Figs. 1B and C). This was detected through GFP signals from mutant mice with a knock-in allele encoding a GFP variant, Venus, fused to the COOH terminus of endogenous FoxO1^32^. The FoxO1-GFP transgene signals were largely colocalized with endogenous FoxO1 protein, as detected by immunofluorescent staining (IF) of FoxO1 (Fig. S1A), validating the specificity of FoxO1-GFP signals. Notably, the FoxO1 signals detected by FoxO1 IF showed consistent decreases in MPOA neurons in females but not males after 2 weeks of HFD feeding (Fig. S1B). These mRNA and protein results suggest that FoxO1 in the MPOA responds to the HFD challenge in a sex-specific manner.

**Figure 1.**
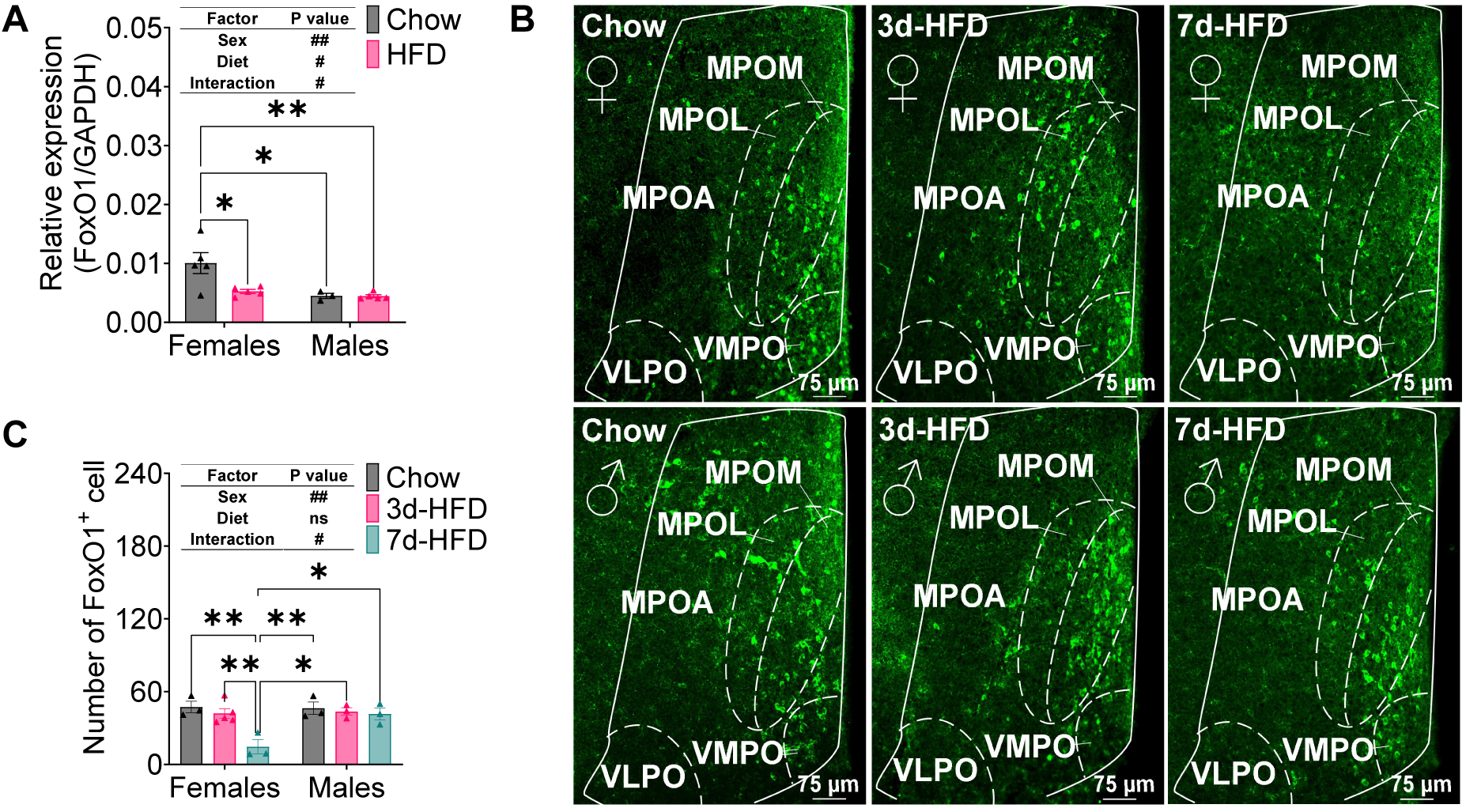
FoxO1^MPOA^ sexual dimorphism responds to a High-Fat Diet (HFD) challenge. ***(A)*** FoxO1 mRNA expression in MPOA after 2-week chow or HFD feeding in male and female C57BL/6J Mice. ***(B-C)*** Representative images (B) and quantification (C) of FoxO1-GFP fluorescence in the MPOA of FoxO1-YFP mice fed either a chow diet, 3-day HFD, or 7-day HFD. Representative images showing the four nuclei of the Medial Preoptic Area (MPOA): Medial Part of Medial Preoptic Nucleus (MPOM), Lateral Part of Medial Preoptic Nucleus (MPOL), Ventral Medial Preoptic Nucleus (VMPO), and Ventrolateral Preoptic Nucleus (VLPO). Results are expressed as means ± SEM. In panels A and C, significance levels are indicated as #P < 0.05 and ##P < 0.01 for two-way ANOVA analysis, and *P < 0.05 and **P < 0.01 for subsequent post hoc Sidak tests.

### Adult FoxO1 deletion in the MPOA affects female fat mass and energy balance

To determine whether alterations in FoxO1^MPOA^ expression affect energy metabolism, we generated a mouse model with FoxO1 selectively deleted in the MPOA (FoxO1-KO^MPOA^) by injecting AAV-CMV-Cre-GFP bilaterally into the MPOA of FoxO1^flox/flox^ mice at 8 weeks of age (Fig. 2A). FoxO1^flox/flox^ littermates received the AAV-CMV-GFP stereotaxic injections and served as controls. Post hoc visualization of GFP and FoxO1 IF confirmed accurate targeting to the MPOA and the efficiency of the deletion in each mouse (Fig. S2A). In female mice under a chow-fed condition, we did not find a difference in body weight between control mice and FoxO1-KO^MPOA^ mice (Fig. 2B). However, FoxO1-KO^MPOA^ significantly increased fat mass and decreased lean mass (Fig. 2C). This was associated with an increased ratio of gonadal white adipose tissue (gWAT) but not brown adipose tissue (BAT) or inguinal white adipose tissue (iWAT, Fig. 2D). These results suggest an obese phenotype characterized by increased fat mass without overall weight gain. Notably, we also found that FoxO1^MPOA^ female mice showed normal glucose homeostasis, indicated by unchanged fed glucose levels, glucose tolerance, and insulin sensitivity (Figs. S2B-D).

**Figure 2.**
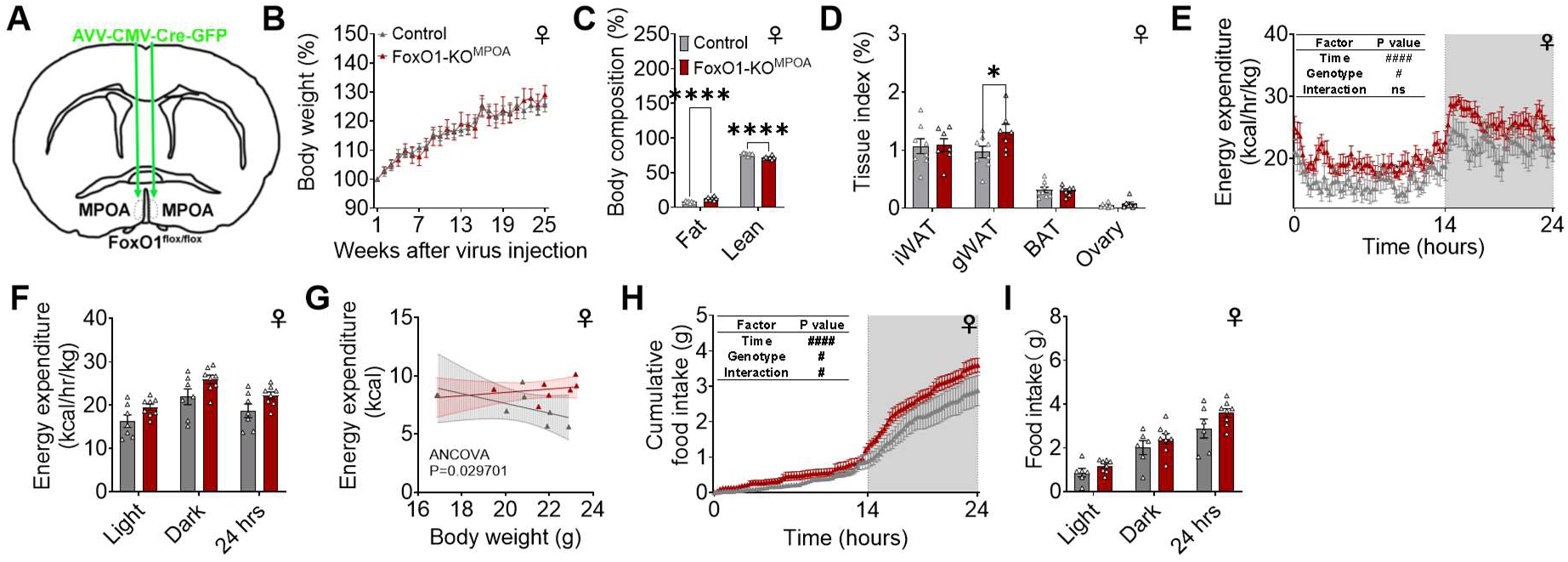
Selective deletion of FOXO1 in the MPOA during adulthood increases fat mass and is associated with higher energy expenditure and food intake in females. ***(A)*** Schematic of the experimental strategy using AAV-CMV-Cre-GFP virus to selectively delete FoxO1 in FoxO1^flox/flox^ mice (FoxO1-KO^MPOA^, 8 weeks). FoxO1^flox/flox^ mice receiving AAV-CMV-GFP virus injections served as controls. ***(B)*** Body weight percentage compared to initial weight at two weeks post-surgery. Female control and FoxO1-KO^MPOA^ mice received virus injections at eight weeks of age and were fed a chow diet for 25 weeks (n = 8/7). ***(C-D)*** Body composition percentage relative to body weight (C) and tissue index percentage relative to body weight (D) in female mice 25 weeks after virus injection (n = 8/7). ***(E-I)*** Energy expenditure (E, n = 7/8), light/dark/24-hour average energy expenditure (F, n = 7/8), ANCOVA analysis of daily total energy expenditure using body weight as a covariate (G, n = 7/8), cumulative food intake (H, n = 6/8), and light/dark/24-hour food intake (I, n = 6/8) in female control and FoxO1-KO^MPOA^ mice. Indirect calorimetry was performed on a separate cohort of mice from those used for chronic body weight recording. These mice had comparable body weight and lean mass at the time of study, 8 weeks after virus injection. Data in panel G were analyzed using ANCOVA with daily total energy expenditure as the dependent variable, genotype as the fixed variable, and body weight as the covariate. Results are displayed as means ± SEM. (E and H) #*P* < 0.05, ####P < 0.0001 in the two-way ANOVA analysis. (C and D) *P < 0.05, ****P < 0.0001 in the two-way ANOVA analysis followed by post hoc Sidak tests.

To determine the mechanisms underlying the obese phenotypes seen in FoxO1-KO^MPOA^ female mice, another cohort of chow-fed FoxO1-KO^MPOA^ females and control littermates (2 months after virus injection at 8 weeks of age), with matched body weight and lean mass, were adapted into the Sable Promethion Core systems to measure energy expenditure and food intake. Female FoxO1-KO^MPOA^ mice showed a modest but significant increase in energy expenditure compared to control mice (Fig. 2E). Although the average energy expenditures during light, dark, and 24-hour periods were not different between FoxO1-KO^MPOA^ and control females (Fig. 2F), ANCOVA analysis of daily total energy expenditure, using body weight as a covariate, indicated a mass-independent difference in energy expenditure, with higher energy expenditure in FoxO1-KO^MPOA^ female mice compared to control female mice (Fig. 2G).

Notably, we did not find any difference in the respiratory quotient (RQ) or physical activity (Figs. S2E-H) induced by FoxO1-KO^MPOA^ in female mice. However, FoxO1-KO^MPOA^ females expressed significantly higher levels of the thermogenic gene uncoupling protein 1 (UCP1) in the BAT, but not in the iWAT or gWAT (Figs. S2I-K). This aligns with previous reports that MPOA neurons regulate energy expenditure partly through modulating thermogenesis in the BAT^28–30,33^.

Interestingly, associated with increased energy expenditure, FoxO1-KO^MPOA^ females consumed a modest but significantly higher amount of food compared to control mice (Figs. 2H and I). This might be a compensatory response to the higher expenditure, which may explain the unchanged body weight. Notably, while room temperature (22 °C) is considered thermoneutral for humans, it is mildly cold for mice, especially when singly housed^34^. The increased energy expenditure induced by FoxO1-KO^MPOA^ could be due to enhanced cold-induced metabolic adaptation, leading to compensatory food intake, eventually resulting in more fat storage to combat a cold environment.

### FoxO1-KO^MPOA^ does not modulate glucose or energy homeostasis in male mice

In contrast to female FoxO1-KO^MPOA^ mice, male mutants did not show differences in body weight, body composition, or adiposity (Figs. S3A-C), indicating no obesity when fed chow. Consistent with unchanged body weight and composition, male FoxO1-KO^MPOA^ mice had similar cumulative food intake, energy expenditure, physical activity, and RQ compared to male control mice (Figs. S3D-H). Additionally, male mutants had similar fed glucose levels, glucose tolerance, and insulin sensitivity compared to male controls (Figs. S3I-K), suggesting normal glucose homeostasis. These results suggest that, unlike females, FoxO1-KO^MPOA^ does not affect glucose or energy homeostasis in male mice, clearly demonstrating a sex dimorphism.

### FoxO1-KO^MPOA^ enhances heat production induced by cold in females

To test the role of FoxO1^MPOA^ in the metabolic adaptation to temperature challenges, we acutely exposed mice to 6, 22, 30, or 37 °C for 6 hours inside the Sable Promethion system during a chow-fed condition in the morning (around 9 am) at 22 °C to monitor temperature stress-induced metabolic responses (Fig. 3A). Interestingly, we found that a 6-hour acute cold exposure significantly increased energy expenditure in female FoxO1-KO^MPOA^ mice compared to female control mice (Fig. 3B). Notably, the stimulatory effects of FoxO1-KO^MPOA^ on energy expenditure during the 6-hour exposure were blunted as the acute exposure temperature increased, specifically at 22, 30, and 37 °C (Figs. 3C-E). Consistently, the average energy expenditure during the 6-hour cold exposure was significantly higher in female FoxO1-KO^MPOA^ mice compared to female control mice, but this difference disappeared as the temperatures increased (Fig. 3F). Additionally, there were no differences in cumulative food intake, physical activity, and RQ during 6-hour period between female mutants and controls at all temperature exposure conditions (Figs. S4A-L). Notably, we did not observe any changes in male mutants during the acute exposure at different temperatures (Figs. S4M-V). These results suggest that FoxO1-KO^MPOA^ selectively enhances acute cold-induced metabolic adaptation in female but not in male mice.

**Figure 3.**
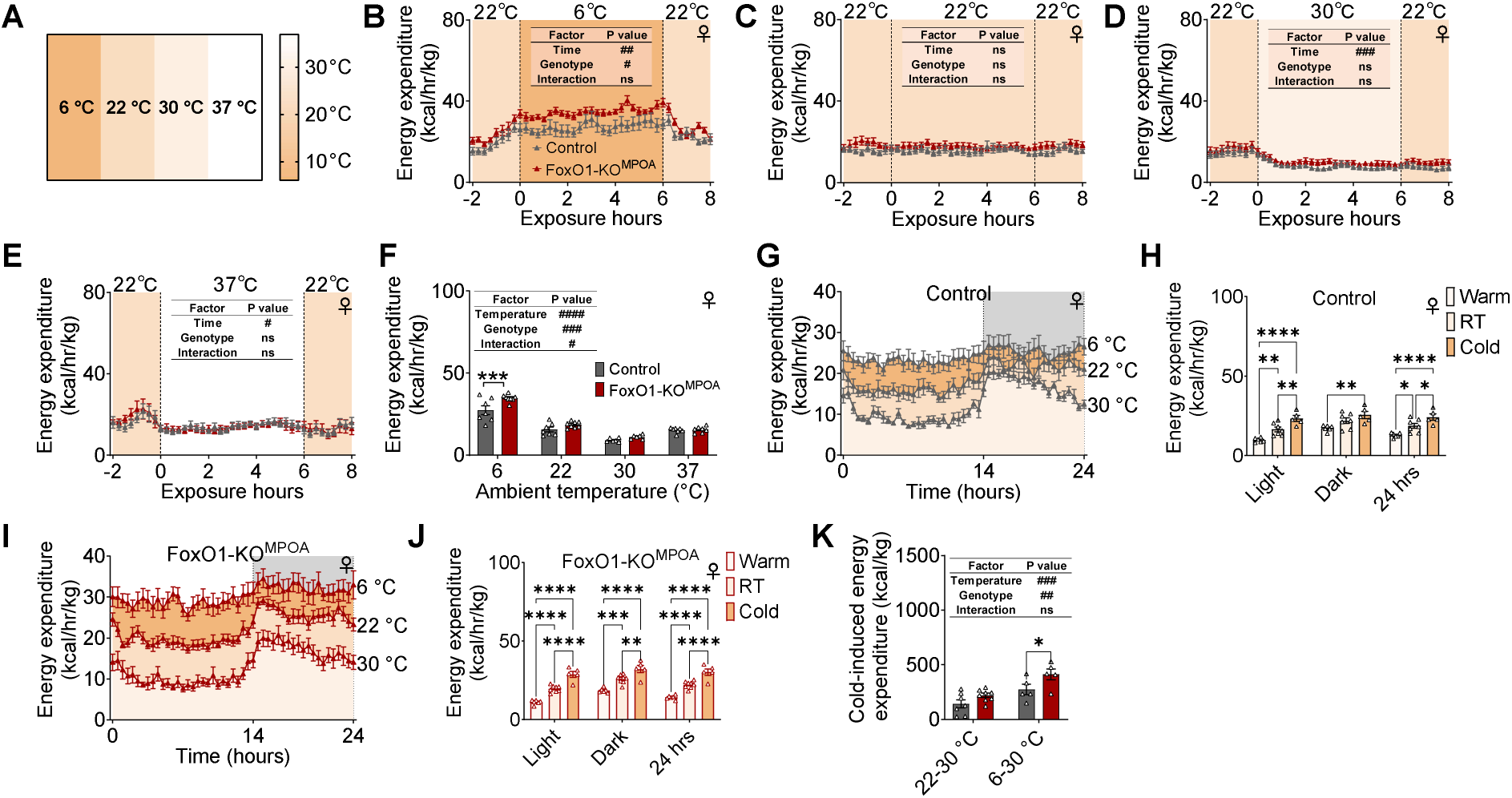
FoxO1-KO^MPOA^ enhances cold-induced heat production in females. ***(A)*** Temperature scheme used for the exposure experiment. ***(B-E)*** Energy expenditure measured before, during, and after six-hour exposure to various temperatures: cold (6°C, B), room temperature (22°C, C), thermoneutrality (30°C, D), and warm (37°C, E, n = 7/8). Mice had comparable body weight and lean mass during the indirect calorimetry study, conducted 8 weeks after virus injection. ***(F)*** Average energy expenditure during temperature exposures (n = 7/8). ***(G-H)*** Energy expenditure (G) and average energy expenditure during light/dark/24 hours (H) in female control mice after 48-hour adaptation to cold, room temperature, and thermoneutrality (n = 5/7/6). Mice had comparable body weight and lean mass during the indirect calorimetry study, conducted 8 weeks after virus injection. ***(I-J)*** Energy expenditure (I) and average energy expenditure during light/dark/24 hours (J) in female FoxO1-KO^MPOA^ mice after 48-hour adaptation to cold, room temperature, and thermoneutrality (n = 5/8/6). ***(K)*** Energy expenditure changes in mice after 48-hour exposure to 22°C or 6°C, compared to thermoneutral conditions at 30°C. Changes calculated from area under the curve from G and I, representing cold-induced heat production (n = 7/8 and 5/5). Results are shown as means ± SEM. (B-F, and K) #*P* < 0.05, ##p < 0.01, ###*P* < 0.001, ####p < 0.0001 in two-way ANOVA. (F, H, J, and K) *p < 0.05, **p < 0.01, \*\*\**P* < 0.001, ****p < 0.0001 in two-way ANOVA followed by post hoc Sidak tests.

To test whether FoxO1^MPOA^ is involved in metabolic adaptation to temperature challenges over a longer period, we adapted female control and FoxO1-KO^MPOA^ mice to thermoneutral (30 °C) or cold (6 °C) environments in the Sable Promethion system for 2 days. During the 48-hour cold exposure, female mutants showed significantly higher energy expenditure and RQ, without changes in cumulative food intake or physical activity (Figs. S5A-G). Conversely, during the 48-hour thermoneutral exposure, the stimulatory effects of FoxO1-KO^MPOA^ on energy expenditure and RQ were diminished in females, with no change in cumulative food intake or physical activity (Figs. S5H-N). These results suggest that the metabolic differences between female control and FoxO1-KO^MPOA^ mice are temperature-dependent, supporting our previous assumption that the increased energy expenditure induced by FoxO1-KO^MPOA^ at room temperature could be due to enhanced cold-induced metabolic adaptation.

Both female control and FoxO1-KO^MPOA^ mice showed temperature-induced energy expenditure adaptation: energy expenditure decreased as the ambient temperature rose and increased as the ambient temperature fell (Figs. 3G-J). However, female mutants had significantly higher heat production triggered by the 6 °C cold exposure compared to control females (Fig. 3K). Additionally, consistent with observations at room temperature (22 °C), male mutants showed no difference in cumulative food intake, energy expenditure, physical activity, and RQ at thermoneutrality (Figs. S5O-U), suggesting a dispensable role of FoxO1^MPOA^ in energy homeostasis regulation and metabolic adaptation to temperature stress. These data support that FoxO1^MPOA^ is essential in the temperature-dependent adaptation of heat production in females but not in males, potentially contributing to the organism’s resilience and homeostasis.

### FoxO1-KO^MPOA^ prevents diet-induced obesity in females

We then characterized metabolic adaptation to nutritional challenges in male and female FoxO1-KO^MPOA^ mice by feeding them a 60% HFD two weeks after virus injection. Similar to the chow-fed condition, male FoxO1-KO^MPOA^ mice showed comparable body weight, fat/lean mass, fat distribution, and food intake to control mice (Figs. S6A–D). Energy expenditure was estimated by evaluating feeding efficiency (the ratio between body weight gain and cumulative food intake), and no changes were observed in male mice (Figs. S6E). In contrast, HFD-fed FoxO1-KO^MPOA^ females showed significant decreases in body weight compared with control females (Fig. 4A). Specifically, HFD-fed FoxO1-KO^MPOA^ females started to be significantly leaner than their controls three weeks after HFD feeding. By 10 weeks after HFD, FoxO1-KO^MPOA^ females gained 5.8 g less body weight than controls (about 21% of control body weight) (Fig. 4A). Fat mass of FoxO1-KO^MPOA^ females decreased by 41%, while lean mass increased by 12% compared to control females (Fig. 4B), both significant. The reduction in fat mass was due to the decreased weight of iWAT, gWAT, and BAT (Fig. 4C). Additionally, the average adipocyte sizes in both iWAT and gWAT were significantly smaller in FoxO1-KO^MPOA^ females than in control females (Figs. 4D-G). These data indicate that FoxO1 in the MPOA of female brains, but not male brains, plays a crucial role in metabolic adaptation to diet-induced obesity.

**Figure 4.**
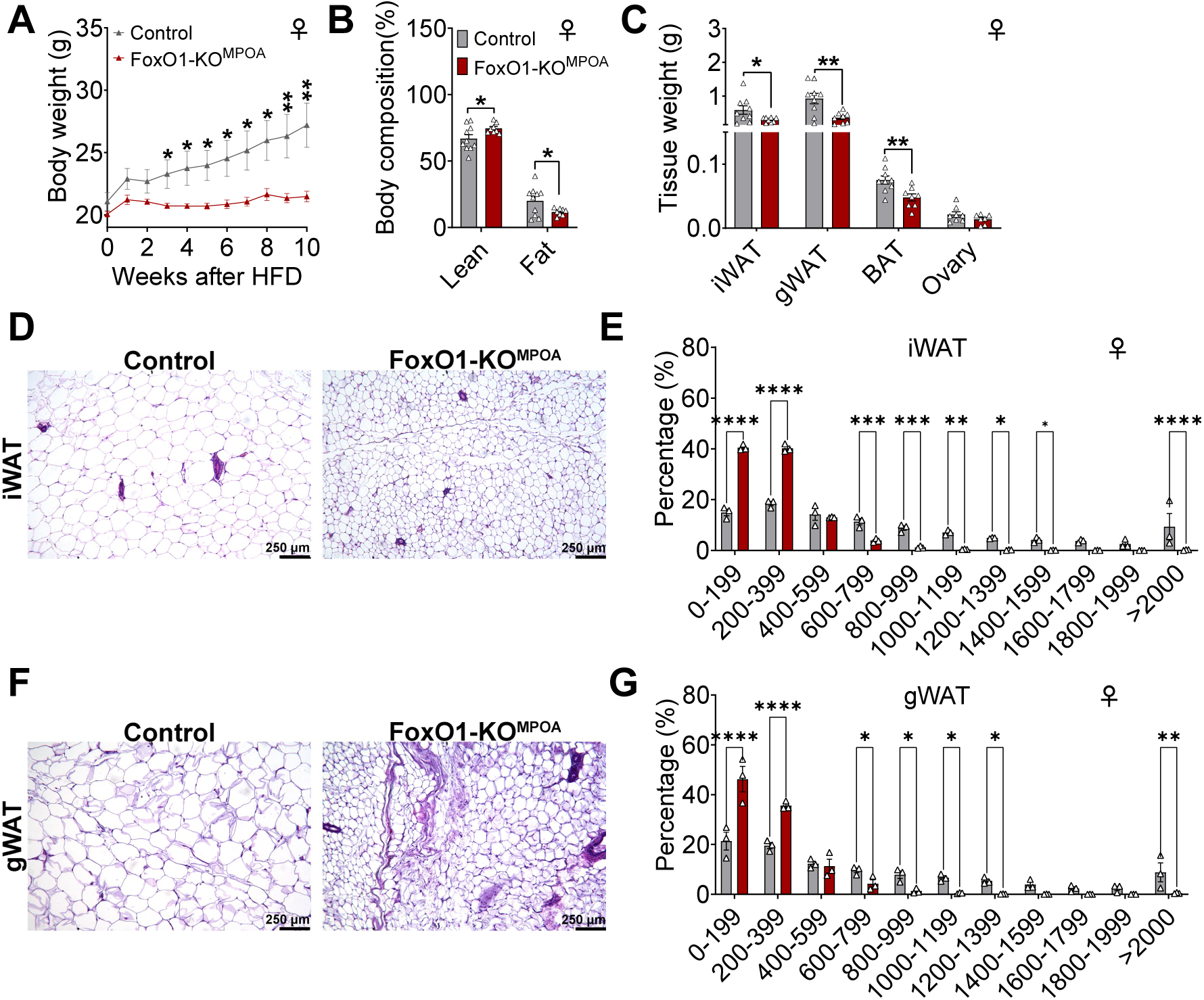
FoxO1-KO^MPOA^ prevents diet-induced obesity in females. ***(A-C)*** Body weight (A), body composition as percentage of total weight (B), and tissue weight (C) of female control and FoxO1-KO^MPOA^ mice after 10 weeks of high-fat diet (HFD) feeding (n = 9/8). HFD feeding began two weeks after virus injection when mice were 10 weeks old. ***(D-G)*** Representative Hematoxylin and Eosin (H&E) staining and cell size quantification in inguinal white adipose tissue (iWAT, D and E) and gonadal white adipose tissue (gWAT, F and G) fat pads after 10 weeks of HFD feeding (n = 3/3). Results are presented as means ± SEM. (A-C, E, and G) \**P* < 0.05, \*\**P* < 0.01, \*\*\**P* < 0.001, \*\*\*\**P* < 0.0001 by two-way ANOVA with post hoc Sidak tests.

Consistent with reduced body weight and adiposity, female mutant mice showed decreased cumulative food intake (Fig. 5A) and feed efficiency (Fig. 5B). This suggests they were less efficient at converting dietary energy into body weight, indirectly indicating increased energy expenditure. These chronic observations were further confirmed by an indirect calorimetry study, revealing reduced overall daily food intake just one day after switching animals from a normal chow diet to a HFD (Figs. 5C-D). Moreover, the energy expenditure of female mutant mice was significantly higher than that of female control mice during both the dark cycle and the 24-hour daily period, just two days after HFD switching (Figs. 5E-F). However, FoxO1-KO^MPOA^ did not affect physical activity in response to an HFD challenge in female mice (Figs. S7A and B). Although the RQ of female mutant mice showed a significant increase during the dark cycle on the first day after the HFD switch, it returned to normal compared to control female mice after two days of diet switching (Figs. S6C and D).

**Figure 5.**
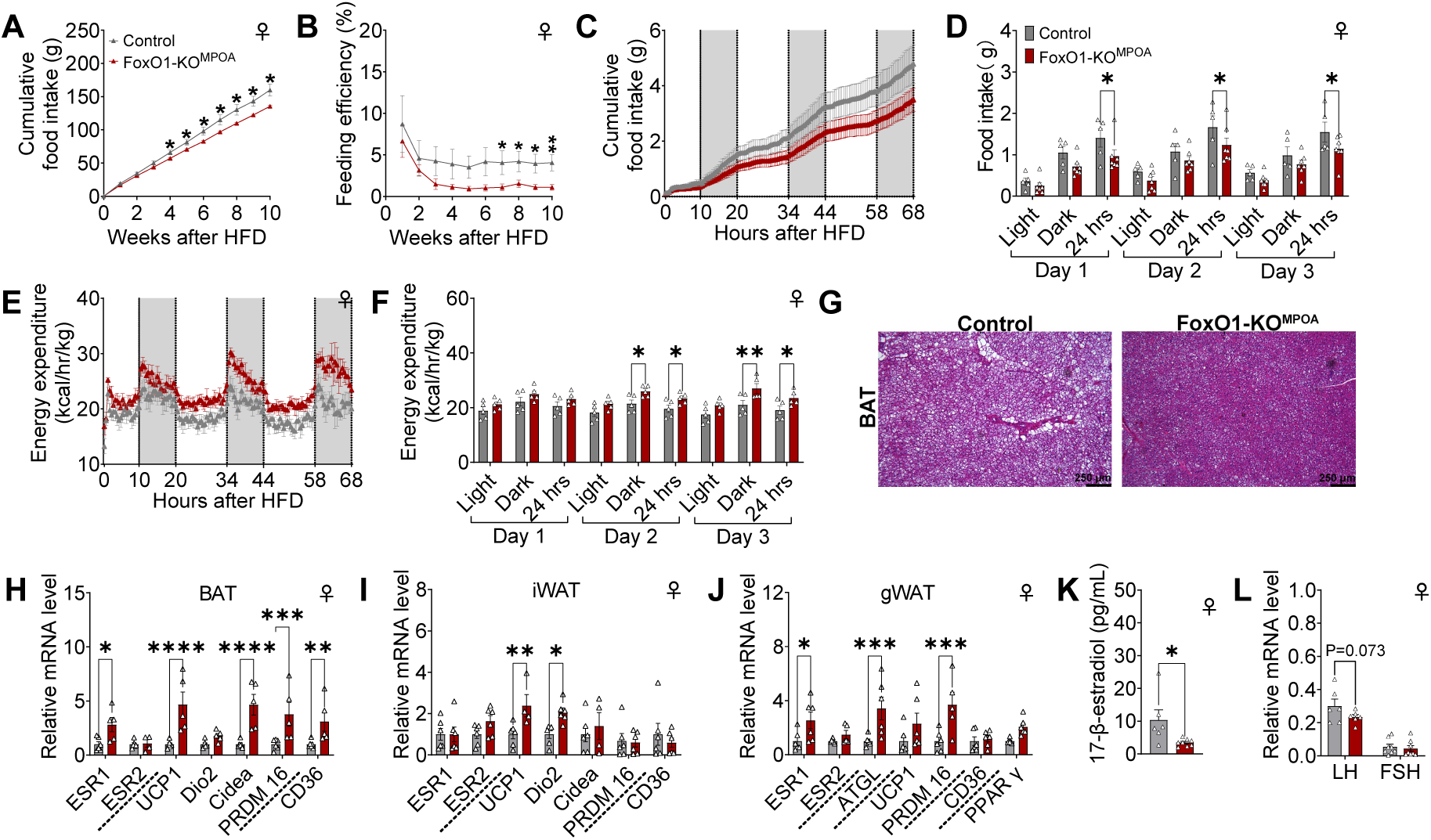
FoxO1-KO^MPOA^ reduces food intake and stimulates energy expenditure during metabolic adaptation to a high-fat diet challenge in female mice. ***(A-B)*** Cumulative food intake (A) and feeding efficiency (B) of female control and FoxO1-KO^MPOA^ mice. Feeding efficiency represents the ratio of body weight change to cumulative food intake (n = 9/8). ***(C-D)*** Cumulative food intake (C) and total food intake during light/dark/24-hour periods (D) of female control and FoxO1-KO^MPOA^ mice following acute adaptation to a high-fat diet challenge (n = 5/6). ***(E-F)*** Energy expenditure (E) and average energy expenditure during light/dark/24-hour periods (F) of female control and FoxO1-KO^MPOA^ mice following acute adaptation to a high-fat diet challenge (n = 5/5). (G) Representative image of brown adipose tissue (BAT) with H&E staining, taken 10 weeks after high-fat diet challenge. ***(H-J)*** mRNA levels of genes in BAT (H), iWAT (I), and gWAT (J) 10 weeks after high-fat diet challenge. Measured genes include estrogen receptors (ESR1 and ESR2), thermogenic genes (UCP1, Dio2, Cidea, PRDM16), lipolytic gene (ATGL), adipogenic gene (PPARγ), and fatty acid sensor and transporter (CD36, n = 6/6). ***(K-L)*** Circulating levels of 17β-estradiol (K, n = 6/8) and relative mRNA levels of LH and FSH to housekeeping gene GAPDH in the pituitary (L, n = 8/8) 10 weeks after high-fat diet challenge. Results are shown as means ± SEM. (A-B, D, and F) \**P* < 0.05, \*\**P* < 0.01 in two-way ANOVA analysis followed by post hoc Sidak tests. (H-K) \**P* < 0.05, \*\**P* < 0.01, \*\*\**P* < 0.001, \*\*\*\**P* < 0.0001 in unpaired *t* tests.

In line with decreased BAT weight, female FoxO1-KO^MPOA^ mice exhibited less lipid deposition compared to female control mice (Fig. 5G). Correspondingly, UCP1 mRNA levels in the BAT and iWAT of female FoxO1-KO^MPOA^ mice were significantly elevated (Fig. 5H and I). Moreover, mRNA levels of Cidea and PR domain containing 16 (PRDM16), factors known to stimulate UCP1 expression^35,36^, and cluster of differentiation 36 (CD36), a fatty acid sensor and transporter crucial for thermogenesis^37^, were significantly higher in female mutant BAT (Fig. 5H). The mRNA expression of iodothyronine deiodinase 2 (Dio2), which encodes an enzyme catalyzing the conversion of T4 to bioactive T3 to stimulate adaptive thermogenesis, was significantly increased in iWAT (Fig. 5I). The mRNA expression of adipose triglyceride lipase (ATGL), a lipolytic gene, and PRDM16 was also upregulated in gWAT (Fig. 5J). Overall, the transcriptional dynamics of adipose tissue in female FoxO1-KO^MPOA^ mice align well with the energy expenditure observations, both suggesting increased thermogenesis and systemic heat production.

These findings suggest a crucial role of FoxO1^MPOA^ in metabolic adaptation to acute and chronic HFD challenges. This adaptation potentially promotes females to increase food intake and reduce energy expenditure, facilitating anabolic pathways and fat storage when energy dense food is available. Such a process may be essential for females to maintain adequate energy reserves for future reproduction or to strengthen metabolic defenses against potential famine. This could represent a conserved evolutionary strategy to enhance female survival and reproductive success.

Interestingly, the levels of ESR1, the gene encoding estrogen receptor α (ERα), were also increased in the BAT and gWAT of female mutant mice (Figs. 5H and J). This was associated with reduced serum 17β-estradiol (E2, Fig. 5K) and mRNA expression of luteinizing hormone (LH) in the pituitary, suggesting potential disruption of estrogen production. This disruption results in lower circulating estrogens and compensatory overexpression of ERα in adipose tissue. However, since estrogens are known to reduce body weight by decreasing food intake and increasing expenditure^38^, it is unlikely that the metabolic phenotypes outlined above are due to the decreased levels of circulating estrogen.

### FoxO1-KO^MPOA^ disrupts metabolic adaptation in response to fasting and refeeding in females

To further explore the effects of FoxO1^MPOA^ on metabolic adaptation to nutritional deficiency, we fasted female FoxO1-KO^MPOA^ mice for 18 hours overnight, followed by a refeeding phase, and monitored their metabolic changes using the Sable Promethion system. Compared to female control mice, the FoxO1-KO^MPOA^ mice showed significantly increased energy expenditure during the dark cycle and tended to have higher average energy expenditure throughout the fasting period, with no changes during refeeding (Figs. S8A-C). Interestingly, this increased energy expenditure was associated with a higher average RQ during the dark cycle of the fasting phase, but not at other times during fasting or refeeding (Figs. S8D-F). As observed in normal chow and HFD conditions, physical activity remained unaffected by FoxO1-KO^MPOA^ in females during both fasting and refeeding phases (Fig. S8G). Notably, FoxO1-KO^MPOA^ significantly reduced total fasting-induced food intake within 20 hours after refeeding (Figs. S8H and I). These data strongly suggest that FoxO1-KO^MPOA^ disrupts the energy reserve defense response during fasting and the fasting-induced refeeding response in females. This FoxO1^MPOA^-mediated metabolic adaptation to nutritional deficiency, and the subsequent rebound in food intake when food becomes available, is potentially crucial for enhancing the survival rate of females during food shortages.

### FoxO1^MPOA^ activation exacerbates DIO in both sexes

To further confirm the role of FoxO1 MPOA signals in the development of DIO, we tested whether FoxO1 activation specifically in the MPOA can promote HFD-induced body weight gain. To this end, we generated a mouse model with FoxO1 constitutively activated in the MPOA by bilaterally injecting AAV-CMV-Cre (FoxO1-CA^MPOA^) or AAV-CMV-GFP (control) virus into the MPOA of both male and female Rosa26-LSL-FoxO1AAA mice (Fig. 6A). These Rosa26-LSL-FoxO1AAA knock-in mice allow Cre-dependent overexpression of a hemagglutinin (HA)-tagged human FOXO1AAA mutant and IRES-GFP proteins. In this FOXO1AAA mutant, amino acids at the Akt phosphorylation sites are replaced with alanines, leading to defective nuclear export of FOXO1 and constitutive activation^39^. Post hoc visualization of GFP was used to confirm accurate targeting to the MPOA and efficient overexpression of FoxO1AAA in each mouse (Fig. 6B). After surgery recovery and individual housing with HFD feeding, both male and female FoxO1-CA^MPOA^ mice showed increased body weight and feeding efficiency, although only males exhibited elevated food intake compared to controls (Fig. 6C-H). These results suggest that FoxO1 activation specifically in the MPOA promotes HFD-induced body weight gain.

**Figure 6.**
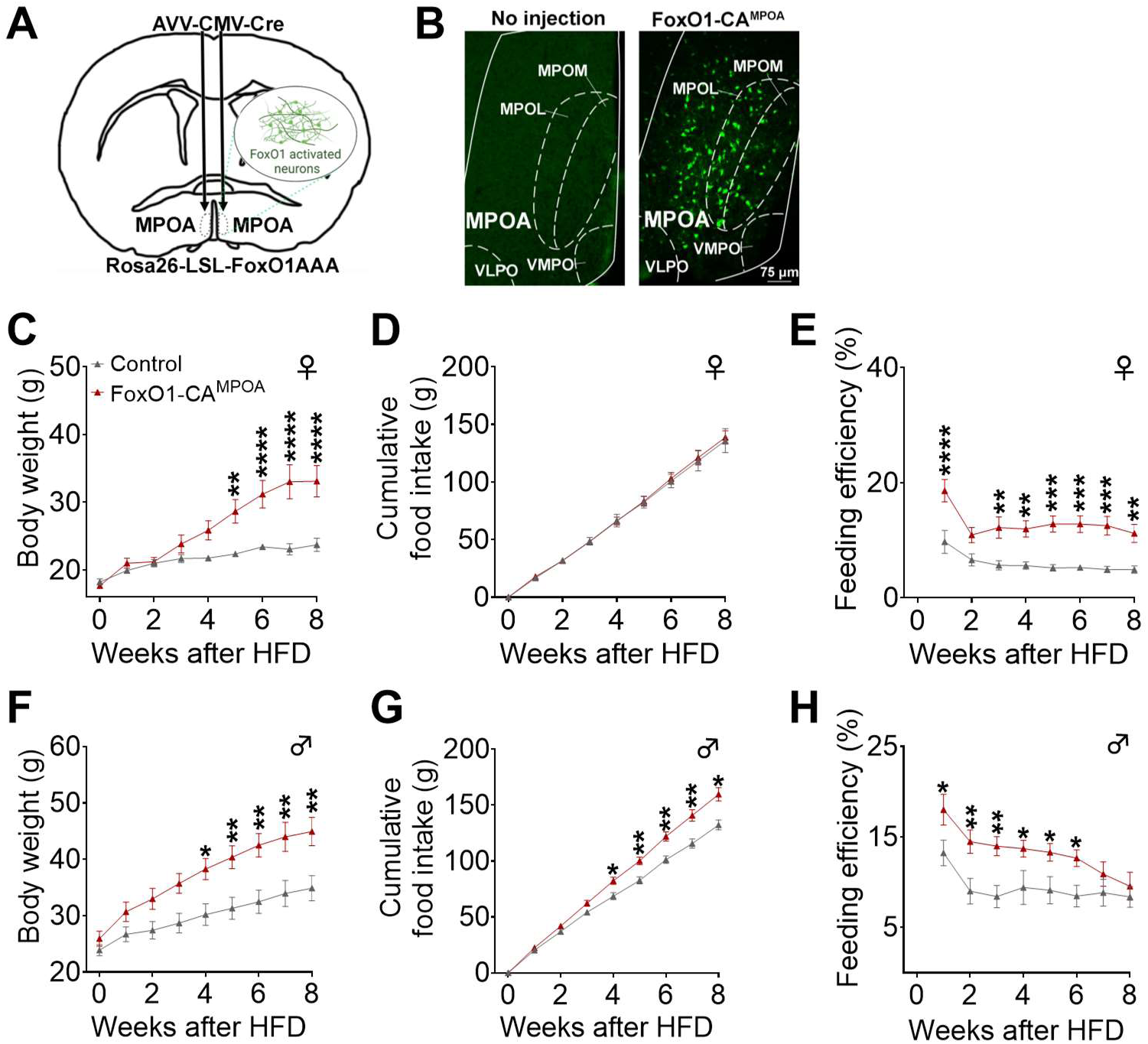
FoxO1-CA^MPOA^ promotes diet-induced obesity in both sexes. ***(A)*** Schematic of the experimental strategy using AAV-CMV-Cre virus to selectively overexpress the constitutively active form of FoxO1AAA/GFP in Rosa26-LSL-FoxO1AAA mice (FoxO1-CA^MPOA^, 8 weeks). Rosa26-LSL-FoxO1AAA mice receiving AAV-CMV-GFP virus injections served as controls. ***(B)*** GFP immunoreactivity was observed in a Rosa26-LSL-FoxO1AAA mouse after AAV-CMV-Cre injection into the MPOA, but not in a Rosa26-LSL-FoxO1AAA mouse without virus injection. ***(C-H)*** Body weight (C and F), cumulative food intake (D and G), and feeding efficiency (E and H) were measured in female and male control and FoxO1-CA^MPOA^ mice. Feeding efficiency was calculated as the ratio of body weight change to cumulative food intake. Mice were fed a high-fat diet for 8 weeks, beginning two weeks after virus injection at 10 weeks of age (females: n = 10/9; males: n = 12/10). Results are shown as means ± SEM. (C-H) \**P* < 0.05, \*\**P* < 0.01, \*\*\**P* < 0.001, \*\*\*\**P* < 0.0001 in two-way ANOVA analysis followed by post hoc Sidak tests.

### The gender-specific anti-DIO effects of FoxO1-KO^MPOA^ require ovarian hormones, but estrogens alone are insufficient

Given that FoxO1-KO^MPOA^ mice showed sex-specific prevention of DIO in females but not males, we investigated whether ovarian hormones played a role in this sex difference. To this end, we generated a cohort of female control and FoxO1-KO^MPOA^ mice that received virus injection combined with OVX at 8 weeks of age. We found that the anti-DIO effect of FoxO1-KO^MPOA^ was completely abolished by ovariectomy and even induced opposite DIO-promoting effects. Specifically, female OVX FoxO1-KO^MPOA^ mice showed a slight, non-significant increase in body weight (Fig. 7A and B). This change was characterized by increased fat mass and decreased lean mass (Fig. 7C), which was attributed to increased gWAT weight (Fig. 7D). These mice consumed more food (Fig. 7E and S9A), while their feeding efficiency, energy expenditure, physical activity, and RQ remained unchanged (Fig. 7F and S8B-D). These findings suggest that the female-specific anti-DIO effects of FoxO1-KO^MPOA^ depend on ovarian hormones.

**Figure 7.**
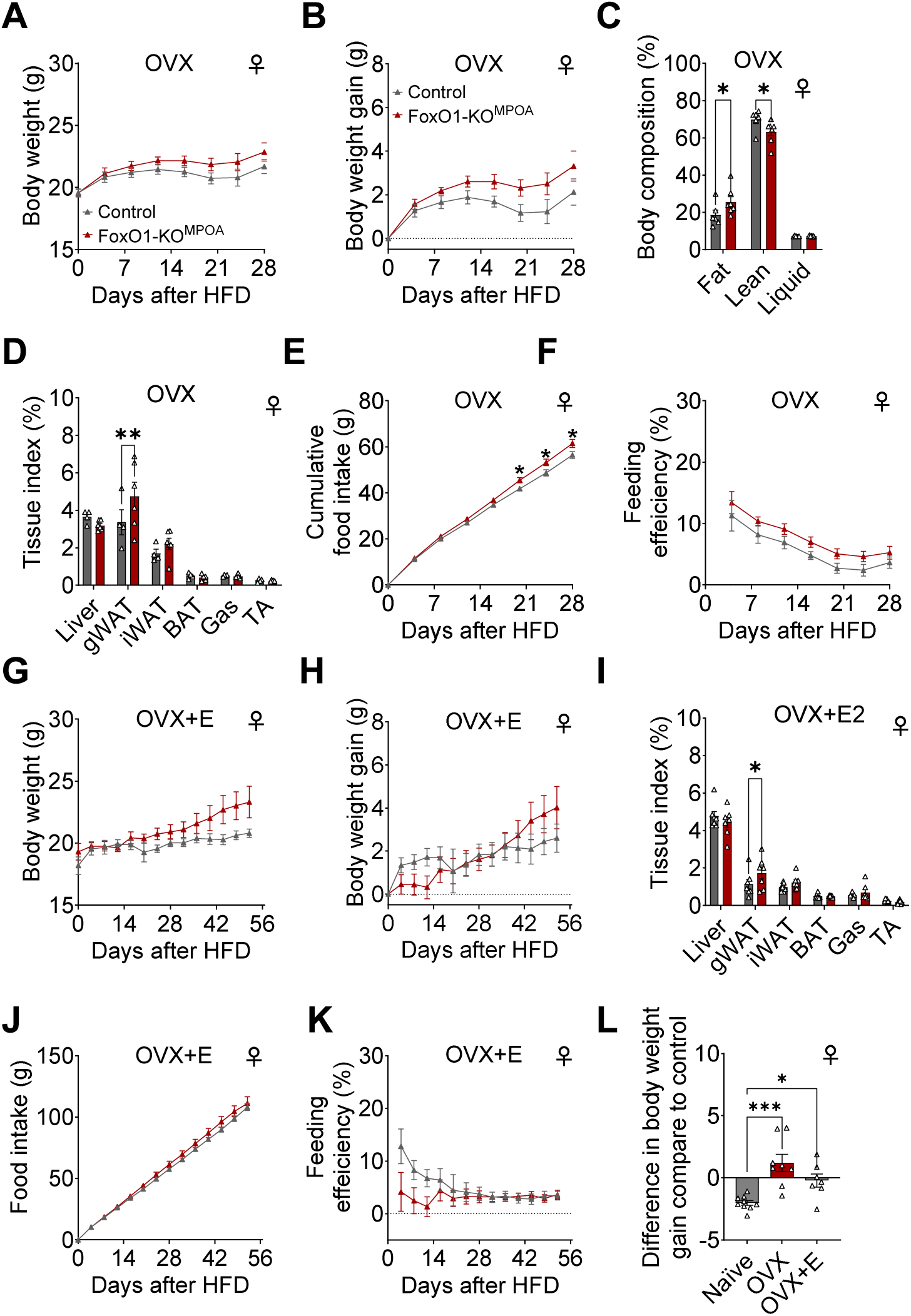
The gender-specific anti-DIO effects of FoxO1-KO^MPOA^ require ovarian hormones, though estrogens alone are not sufficient. ***(A-K)*** Analysis of body weight, body weight gain, body composition, tissue index (as percentage of total weight), cumulative food intake, and feeding efficiency in two groups: ovariectomized female control and FoxO1-KO^MPOA^ mice (OVX, n = 7/8, A-E), and OVX female mice with subcutaneous 17β-estradiol pellet implants (OVX+E, 0.025 mg/pellet, 60-day maximum release, n = 7/7, F-J). All mice received a high-fat diet for 28-56 days, beginning four days after virus injection and either OVX or OVX+E pellet implantation at 8 weeks of age. Gas: gastrocnemius muscle; TA: tibialis anterior muscle. ***(L)*** Comparison of body weight gain between female FoxO1-KO^MPOA^ mice and control mice after 4 weeks of HFD feeding under naïve, OVX, or OVX+E conditions (n = 9/8/7). Results are shown as means ± SEM. (C, D, and I) \**P* < 0.05, \*\**P* < 0.01 in two-way ANOVA followed by post hoc Sidak tests. (L) \**P* < 0.05, \*\*\**P* < 0.001 in one-way ANOVA followed by post hoc Dunnett tests.

Next, we tested whether estrogen supplementation can restore the anti-DIO effects of FoxO1-KO^MPOA^. Specifically, we administered 17β-estradiol pellets (0.025 mg/p, 60-day maximum release) subcutaneously to the ovariectomized female control and FoxO1-KO^MPOA^ mice. We found that estrogen supplementation following ovariectomy did not restore the anti-DIO effects of FoxO1-KO^MPOA^ (Fig. 7G-L), indicating that estrogen alone is not sufficient to produce these sex-specific effects. In line with these findings, when we supplemented male control and FoxO1-KO^MPOA^ mice with 17β-estradiol pellets, we observed no changes in body weight, food intake, feeding efficiency, energy expenditure, physical activity, or RQ (Fig. S8E-K). These results confirm that while the female-specific anti-DIO effects of FoxO1-KO^MPOA^ require ovarian hormones, estrogen alone cannot reproduce these effects.

To investigate potential molecular mechanisms behind sex-specific metabolic adaptation in the MPOA, we reanalyzed published single-cell RNA sequencing data from the MPOA^40^. Using t-distributed stochastic neighbor embedding (tSNE) dimensionality reduction, we clustered cells into 13 distinct groups and annotated them using SingleR and Azimuth (Fig. S10A). Our analysis revealed FoxO1 expression predominantly in inhibitory neurons (Fig. S10B and C). FoxO1 co-localizes with steroid hormone receptors, including ESR1, estrogen receptor 2 (ESR2), G protein-coupled estrogen receptor 1 (GPER1), progesterone receptor (PR), and androgen receptor (AR), in these inhibitory neurons in both male and female mice (Fig. S10D). Within FoxO1-positive inhibitory neurons, PR and AR demonstrated the highest co-expression rates, ESR1 showed moderate co-expression, and ESR2 and GPER1 exhibited minimal co-expression (Fig. S10E). These findings suggest a potential regulatory network between FoxO1 and sex hormone signaling, specifically through PR pathways, in female MPOA inhibitory neurons.

## Discussion

Our findings reveal a previously unrecognized, sex-specific role of FoxO1^MPOA^ in mediating metabolic adaptation to both nutritional and temperature stresses. While FoxO1-KO^MPOA^ did not affect body weight in mice on a normal chow diet, it conferred robust protection against DIO in female mice by increasing energy expenditure, enhancing thermogenic activity, and reducing food intake. The stimulatory effects on energy expenditure were temperature-dependent, emerging most pronounced during cold exposure. The anti-DIO effects of FOXO1-KO^MPOA^ were abolished by ovariectomy and were not rescued by estradiol supplementation, suggesting that ovarian hormones are required but estrogen alone is insufficient. The importance of FOXO1^MPOA^ in metabolic adaptation was further enhanced by the observations that the constitutive activation of FOXO1^MPOA^ promoted DIO in both sexes, highlighting a broader role for FoxO1 in driving energy storage during nutrient excess.

FoxO1 plays a crucial role in regulating energy homeostasis through multiple mechanisms across brain regions. When deleted from most hypothalamic neurons, it leads to reduced body weight and fat mass in both sexes^41^. Its metabolic effects vary depending on the neuronal subset. In the arcuate nucleus (ARC), FoxO1 increases food intake and adiposity by promoting AgRP while inhibiting POMC expression in male and female mice^16,18,19^. When FoxO1 is deleted from either ventromedial hypothalamus (VMH) SF-1-positive neurons^20^ or mesolimbic dopamine neurons^21^, body weight decreases through increased energy expenditure and enhanced BAT thermogenesis, while food intake remains unchanged in both sexes. Although these studies demonstrate a critical role of FoxO1 in energy homeostasis regulation in both males and females, its potential function in metabolic stress resistance and sex-specific adaptation remains unknown. In this study, we systematically examined how FoxO1 in the MPOA, a key integrating center for the regulation of temperature and energy homeostasis^42^, affects adaptation to metabolic stresses (both nutritional and temperature-related). We found that MPOA^FoxO1^ is crucial for metabolic stress adaptation in females but not in males. This finding suggests the existence of a female-specific ancestral stress-resistance pathway, likely evolved to support females cope with the energetic demands of gestation and lactation.

Our findings demonstrate that FoxO1^MPOA^ modulates food intake and energy expenditure in a temperature- and nutritional status-dependent manner. This aligns with previous observations that MPOA neurons maintain body temperature in response to temperature challenges by adjusting thermogenesis, which in turn drives compensatory changes in food intake to maintain energy homeostasis during temperature adaptation^24–27^. Specifically, temperature-sensitive MPOA neurons maintain temperature homeostasis by modifying BAT thermogenesis and energy expenditure through the sympathetic nervous system (SNS) in response to ambient temperature changes, increasing during cold exposure and decreasing during warm exposure^24,25^. These neurons correspondingly balance energy expenditure by coordinating food consumption, preventing weight loss in cold conditions and weight gain in warm conditions^24,26,27^. Through this interplay between energy expenditure and food intake, MPOA maintains both temperature and energy homeostasis. Recent optogenetic and neuronal tracing studies have identified two main temperature-dependent neural circuits in the MPOA that control feeding behavior. The ARC-projecting glutamatergic neurons in MPOA sense low temperature and promote food intake, while the paraventricular hypothalamic nucleus (PVH)-projecting glutamatergic neurons in MPOA are primarily sensitive to high temperature and suppress food intake^43,44^. These temperature- and circuit-specific functions align with our finding that FoxO1^MPOA^ regulates food intake and energy expenditure based on both temperature and nutritional status. However, further research is needed to understand the precise circuit mechanisms underlying these FoxO1^MPOA^ functions.

FoxO1-KO^MPOA^ did not affect body weight in either sex when fed a chow diet at room temperature. In female mice, however, it altered body composition by increasing fat proportions while decreasing lean mass, associated with elevated 24-hour food intake, energy expenditure, and BAT thermogenesis. These parallel increases in energy intake and expenditure maintained overall energy balance, keeping body weight stable. The altered fat/lean ratio in female FoxO1-KO^MPOA^ mice, most reflected by increased gWAT tissue percentage, showed redistribution of energy storage. This shift may represent a compensatory mechanism to store more energy substrates needed for increased heat production. Although increased food intake could contribute to higher energy expenditure through diet-induced thermogenesis, female FoxO1-KO^MPOA^ mice maintained elevated energy expenditure even during fasting, indicating a feeding-independent mechanism. These findings suggest that at room temperature (a mild cold environment for mice), FoxO1-KO^MPOA^ drives increased energy expenditure and BAT thermogenesis, leading to compensatory hyperphagia to offset heat-related energy loss.

Our study revealed a novel temperature-dependent phenotype in female FoxO1-KO^MPOA^ mice that significantly impacts energy expenditure regulation, particularly during cold exposure. Through both acute (6-hour) and extended (48-hour) exposure experiments, we observed that FoxO1-KO^MPOA^ mice exhibited a significant increase in energy expenditure at cold temperatures (6°C), with diminishing effects at room temperature (22°C) and no differences at thermoneutral (30°C) or warm (37°C) conditions. This temperature-dependent response suggests that FoxO1 in MPOA neurons functions as a regulatory brake on cold-induced thermogenic responses, likely through modulation of sympathetic outflow to thermogenic tissues or regulation of local thermal sensing circuits. The specific activation during cold exposure, without effects at warmer temperatures, indicates that FoxO1 primarily regulates cold-defensive rather than heat-defensive pathways in MPOA neurons.

We further demonstrate that cold-induced energy expenditure is enhanced in female FoxO1-KO^MPOA^ mice, but not males, suggesting a sex-specific function for FoxO1^MPOA^ in thermoregulation. This aligns with established research showing that females typically display higher cold-induced thermogenesis than males, attributed to their higher BAT activity, enhanced thermogenic responses mediated by estrogen, and distinct shivering thresholds^45–49^. The molecular mechanism potentially involves estrogen signaling through two pathways: either estrogen receptor α (ERα) directly interacts with FoxO1 to suppress FoxO1-mediated transcription^50^, or membrane-bound ERα activates the PI3K-Akt pathway to phosphorylate and inhibit FoxO1^51–53^. Supporting this hypothesis, we found that FoxO1 and ESR1 (ERα) are co-expressed in MPOA neurons. However, additional research is needed to fully understand how FoxO1 mediates sex-specific differences in cold-induced thermogenesis and whether this occurs through the estrogen/ERα/PI3K/Akt signaling pathway. If this holds true, this interaction may explain why cold-induced thermogenesis varies throughout the menstrual cycle in humans, particularly during the follicular phase when estrogen levels are highest^45,54,55^.

Another important finding is that FoxO1-KO^MPOA^ exhibits strong anti-obesity effects during HFD feeding in female but not male mice. After 10 weeks of HFD feeding, female FoxO1-KO^MPOA^ mice showed significant reductions in body weight through decreased fat mass (including iWAT, gWAT, and BAT) and increased lean mass, accompanied by reduced adipose tissue size, enhanced BAT thermogenesis, and white adipose tissue browning. These metabolic changes were coupled with increased energy expenditure and decreased food intake immediately following the switch from chow to HFD. This response pattern differs from the temperature challenge response during chow-fed conditions, which primarily affects cold-induced defensive thermogenesis without impacting acute food intake.

Supporting these findings, the opposite phenotype was observed in FoxO-CA^MPOA^ mice, which showed significant increases in HFD-induced body weight gain and feeding efficiency, indicating decreased energy expenditure in both sexes. While female FoxO-CA^MPOA^ mice maintained normal food intake, males showed increased food consumption 4 weeks after significant body weight gain, suggesting that reduced energy expenditure is the primary driver of body weight changes under HFD conditions in both sexes. The contrasting effects between FoxO1-KO^MPOA^ and FoxO-CA^MPOA^ on body weight and energy expenditure, but not food intake, may be attributed to FoxO-CA’s constitutive expression in all MPOA Cre-positive cells, including those that normally do not express FoxO1.

The female-specific effects of FoxO1^MPOA^ on DIO suggest an evolutionarily conserved mechanism for energy storage in females. This sex-dependent response likely evolved as a protective adaptation to ensure survival during food scarcity and support the high energy demands of gestation and lactation. Studies have revealed a strong evolutionary connection between sex chromosomes, hormones, and energy balance, potentially explaining females’ enhanced resistance to famine and their greater capacity to store fat when food is abundant^56,57^. Research further demonstrates that women exhibit superior energy conservation and fat storage abilities, especially during pregnancy. Women develop a higher percentage of fat mass in early pregnancy without significantly increasing energy intake, revealing specific metabolic adaptations that drive sex-based differences in fat mass regulation^58,59^. Thus, the pronounced anti-obesity effects seen in female FoxO1-KO^MPOA^ mice during HFD feeding likely represent the disruption of this female-specific energy conservation mechanism, underscoring FoxO1^MPOA^’s crucial role in sex-specific metabolic adaptation to an energy surplus condition.

Since sex hormones contribute to females’ higher capacity to store fat during an energy surplus condition^56,57^, we investigated whether ovarian hormones regulate FoxO1 to mediate sex-specific effects on DIO. We found that depleting endogenous ovarian hormones blocked the anti-obesity effects of FoxO1-KO^MPOA^ on both body weight and food intake during HFD feeding, suggesting that ovarian hormones are necessary for the sex-specific anti-DIO effects of FoxO1-KO^MPOA^. Initially, we hypothesized that estrogen might be the key hormone acting as an upstream modulator of the FoxO1 signaling pathway. However, when we attempted to restore these effects through E2 supplementation, E2 pellets neither restored the anti-obesity effects in OVX FoxO1-KO^MPOA^ females nor produced anti-DIO effects in naive FoxO1-KO^MPOA^ males. Moreover, previous studies have shown that estrogens prevent rather than promote DIO^60–62^ and inhibit rather than activate FoxO1 signaling^51–53^. While estrogen and FoxO1 interaction may play a role in cold-induced thermogenesis, these findings suggest that estrogen is unlikely to be the primary ovarian hormone interacting with FoxO1 to promote energy storage during energy surplus conditions.

Instead, evidence points to progesterone as a more likely upstream modulator for interaction with FoxO1^MPOA^ in regulating energy balance during HFD feeding in females, supported by several findings. The MPOA exhibits abundant expression of progesterone receptors^63^, and research has demonstrated that progesterone receptors not only activate FoxO1 but also trigger FoxO1-dependent signaling pathways in ovarian cells^64^. Furthermore, multiple rodent studies have established that progesterone promotes DIO by stimulating food intake^65–68^. Our own observations strengthen this hypothesis, as we found higher colocalization between FoxO1^MPOA^ positive neurons and progesterone compared to ERα. While these findings collectively suggest that progesterone-FoxO1 interactions may be the potential mechanism through which progesterone influences energy balance, additional research is needed to test this hypothesis directly.

In summary, this study identifies FoxO1 in MPOA neurons as a critical regulator of sex-specific metabolic adaptation, particular in females. Our findings reveal a previously unknown central neural mechanism governing energy balance that integrates temperature, nutrition, and hormonal signals. These results not only advance our understanding of metabolic regulation but also highlight potential therapeutic strategies for obesity and metabolic disorders through sex-specific, brain-targeted FoxO1 modulation.

## Methods

### Mice

All mice were maintained on a C57BL6/J background. The mouse lines included Foxo1^flox/flox^ (#024756, Jackson Laboratory, Bar Harbor, ME), Rosa26-LSL-FoxO1AAA (#029316, Jackson Laboratory), and FoxO1-GFP (obtained from Dr. Domenico Accili at Columbia University)^32^. Mice were housed in a temperature-controlled environment at 22-24°C on a 14-hour light/10-hour dark cycle (6 am to 8 pm). Unless otherwise stated, mice were fed *ad libitum* with a standard mouse chow (6.5% fat; #2920, Harlan-Teklad, Madison, WI) and water. High-fat diet (60% fat; #D12492i) was purchased from Research Diets (New Brunswick, NJ). Animal care and procedures were approved by the Institutional Animal Care and Use Committees at The University of Illinois Chicago.

### FoxO1 expression in response to nutritional challenges

FoxO1-GFP transgenic mice were used, which express Venus (a GFP variant) fused to the COOH terminus of the endogenous FoxO1 protein. At 8 weeks of age, mice were randomly assigned to receive either a standard chow diet or HFD for 3 or 7 days. Following cardiac perfusion, brain tissue was collected and sectioned into 30-μm slices. For immunofluorescence staining, sections were blocked in 3% normal donkey serum for 1 h at room temperature, followed by overnight incubation at room temperature with chicken anti-GFP antibody (1:1000, GFP-1020, AVES Labs) under continuous agitation. Sections were then incubated with biotin-SP-conjugated donkey anti-chicken secondary antibody (1:1000, 703-065-155, Jackson ImmunoResearch) for 2 h and subsequently with Alexa Fluor 488-conjugated streptavidin (1:500, 016-540-084, Jackson ImmunoResearch) for 1 h. Sections were mounted using anti-fade mounting medium (H-1500, Vector Laboratories). Images were acquired using a Leica DM5500 fluorescence microscope. Quantification of GFP-positive neurons was performed on five consecutive coronal sections containing the MPOA nucleus per mouse (n = 3 mice per group), with the average count per mouse representing one biological replicate.

For endogenous FoxO1 expression analysis, C57BL/6J mice were fed HFD for two weeks and processed following the same immunostaining protocol, except that rabbit monoclonal anti-FoxO1 antibody (1:1000, C29H4, 2880S, Cell Signaling Technology) and goat anti-rabbit Alexa Fluor 488-conjugated secondary antibody (1:500, 111-545-003, Jackson ImmunoResearch) were used as primary and secondary antibodies, respectively. FoxO1 mRNA expression was analyzed by RT-PCR using punch-out MPOA tissue from a separate cohort of C57BL/6J mice, following published protocols^69^. All primer sequences are provided in Table 1. Expression data were normalized to *Gapdh* as internal control.

**Table 1.**
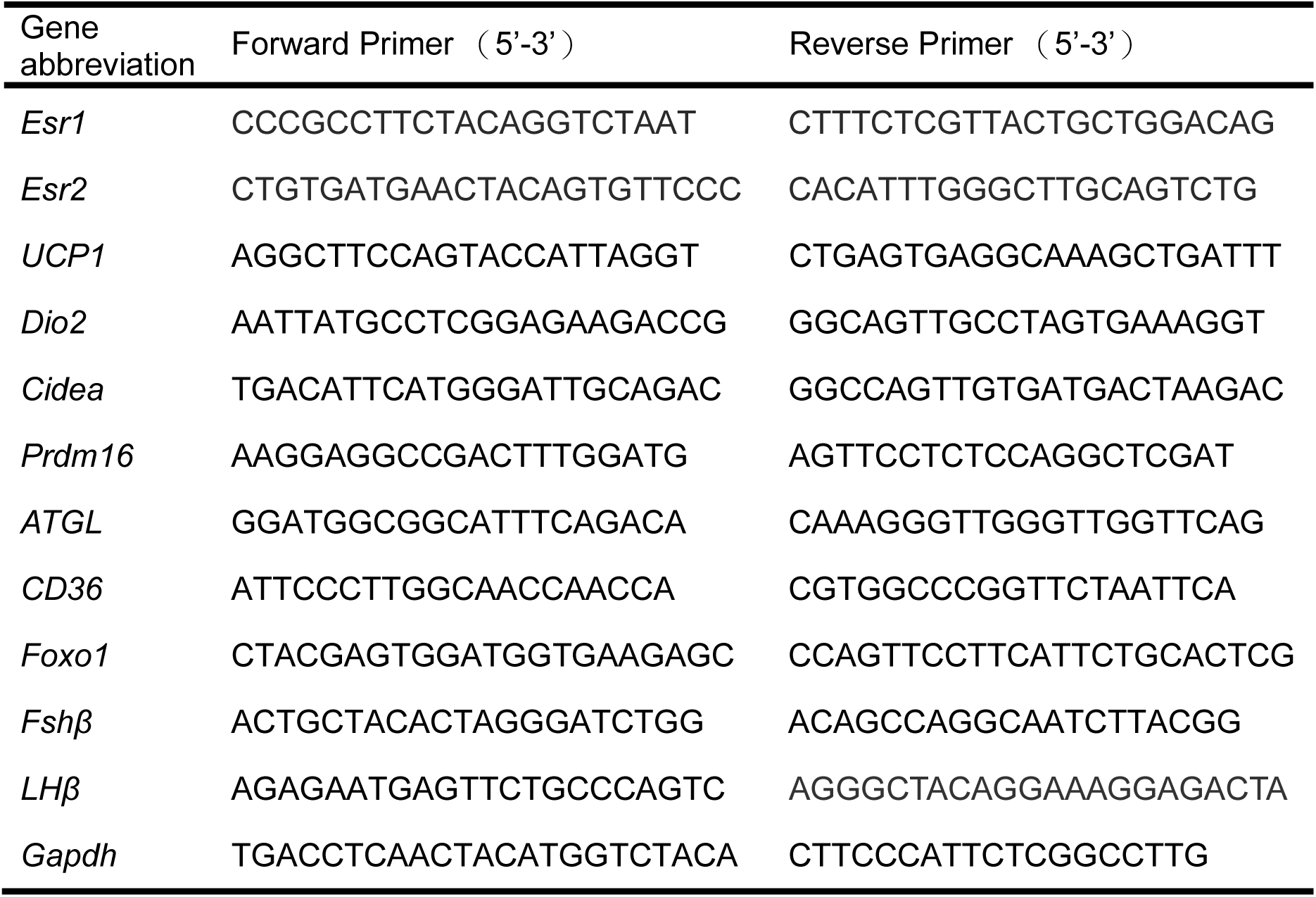
qPCR primer sequences of related genes.

To validate the correlation between FoxO1-GFP transgene and endogenous FoxO1 protein expression, co-immunostaining was performed on 30-μm MPOA brain sections from chow-fed male FoxO1-GFP mice. Following GFP detection as described above, sections were incubated with rabbit anti-FoxO1 primary antibody overnight and goat anti-rabbit Alexa Fluor 594-conjugated secondary antibody for 2 h. Sections were mounted with anti-fade medium and visualized using a Leica DM5500 microscope. FoxO1-positive neurons were detected through GFP-derived signals (Alexa Fluor 488; green) and antibody-based FoxO1 labeling (Alexa Fluor 594; red).

### Adulthood FoxO1 MPOA deletion

To investigate the metabolic role of FoxO1^MPOA^, we generated MPOA-specific FoxO1 knockout mice using a virus-mediated Cre-loxP system. Adult FoxO1^flox/flox^ mice (8 weeks old) received bilateral stereotaxic injections of either AAV-CMV-Cre-GFP or AAV-CMV-GFP (UNC Vector Core; 4×10^12^ GC/mL; 200 nL per side) into the MPOA (bregma, anterior-posterior: 0.26 mm; central-lateral: ±0.35 mm; dorsal-ventral: −5.2 mm) to generate knockout and control groups, respectively.

Metabolic phenotypes were assessed through four experimental paradigms. First, body weight was monitored in group-housed mice maintained on a standard chow diet for 25 weeks. Second, metabolic adaptation to temperature challenges was analyzed using indirect calorimetry in singly-housed, chow-fed mice. Third, metabolic response to HFD was assessed in singly-housed mice by tracking food intake and body weight over 10 weeks. Fourth, acute metabolic adaptation was evaluated during fasting-refeeding and diet transition from a standard chow to HFD using indirect calorimetry. Successful MPOA targeting and FoxO1 deletion were validated via post-mortem analysis of GFP expression and FoxO1 immunofluorescence in all experimental animals. Terminal adipose tissues were collected for qPCR or H&E staining. Serum was collected to measure 17β-estradiol using gas chromatography-mass spectrometry (GC-MS) as previously described^70,71^.

### Indirect calorimetry

Indirect calorimetry measurements were conducted using the Promethion metabolic cage system (Sable Systems International, Las Vegas, NV, USA) at the Metabolic Phenotyping Core, Biologic Resources Laboratory, University of Illinois Chicago. Mice underwent a 72-hour acclimatization period before experiments, with unrestricted access to standard mouse chow and water. All instrument control and data acquisition procedures adhered to manufacturer’s guidelines^72^.

To assess the role of FoxO1 in MPOA-mediated metabolic adaptation to temperature challenges, control and FoxO1-KO^MPOA^ mice were subjected to varying ambient temperatures in the Promethion system. For acute thermal challenges, ambient temperature was maintained at 6°C for 6 hours (9 am to 3 pm) with hourly monitoring. Following a 24-hour recovery period at room temperature (22°C), the procedure was repeated at 30°C and 37°C. For prolonged temperature challenges, a separate cohort of mice was maintained at either 30°C or 6°C for 48 hours following the acclimatization period.

To investigate the role of FoxO1 in MPOA-mediated metabolic adaptation to nutritional deficiency, control and FoxO1-KO^MPOA^ mice maintained on a standard chow underwent an 18-hour fasting period (7 pm to 1 pm), followed by refeeding. Water was provided *ad libitum* throughout the experiment. Following a 7-day recovery period, the role of FoxO1 in MPOA-mediated metabolic adaptation to overnutrition was assessed by monitoring metabolic parameters for 72 hours after transitioning from a standard chow to HFD. Standard chow was replaced with HFD at 9 am, and metabolic measurements were recorded continuously throughout the experimental period.

Indirect calorimetry data were analyzed using ExpeData software (Sable Systems) and CalR (https://calrapp.org). Energy expenditure analysis was performed using ANCOVA through the NIDDK Mouse Metabolic Phenotyping Centers (MMPC) Energy Expenditure Analysis platform (http://www.mmpc.org/shared/regression.aspx).

### RNA isolation and qPCR

To assess MPOA FoxO1 mRNA expression levels during the nutritional challenge, MPOA punches (0.8 mm diameter) were collected from wild-type mice fed either a chow diet or a HFD for 2 weeks, according to established protocols^73^. To assess thermogenesis and fatty acid metabolism in FoxO1-KO^MPOA^ mice, samples were collected from BAT, iWAT, and gWAT. RNA was isolated and purified using TRIzol (#15596018, Invitrogen, Carlsbad, CA) in combination with the Aurum™ Total RNA Fatty and Fibrous Tissue Kit (#7326830, Bio-Rad, Hercules, CA). Quantitative PCR (qPCR) analysis was performed using 2X Universal SYBR Green Fast qPCR Mix (#RK21203, ABclonal, Woburn, MA) according to the manufacturer’s protocol. All primer sequences are provided in Table 1. Expression data were normalized to *Gapdh* as internal control.

### H&E staining

Histological differences in adipose tissue between control and FoxO1-KO^MPOA^ mice were assessed using Hematoxylin and Eosin (H&E) staining of BAT, iWAT, and gWAT. Tissue samples were fixed in 10% neutral buffered formalin at room temperature for 24-48 hours, followed by storage in 70% ethanol. The fixed tissues were paraffin-embedded, sectioned at 5 μm thickness, and stained with H&E. Images were acquired using a Leica DM5500 microscope. Adipocyte size measurements in iWAT and gWAT were performed using ImageJ software. For each mouse, measurements were averaged across three different tissue sections, with each averaged value representing one biological replicate. Statistical analyses were conducted using data from three biological replicates (n=3 mice per group).

### Glucose homeostasis tests

Glucose tolerance test (GTT) was performed to evaluate the role of FoxO1^MPOA^ in glucose metabolism. Mice were fasted overnight (13 h) from 7 pm with *libitum* access to water. Following the fast, baseline blood glucose measurements were obtained via tail vein sampling at 8 am using a StatStrip Xpress 2 Glucose Meter (56510, Nova Biomedical, Waltham, MA). Subsequently, mice received an intraperitoneal injection of dextrose (D9434, Sigma-Aldrich; 2 g/kg body weight). Blood glucose levels were monitored at 15, 30, 60, and 120 minutes post-injection to assess glucose clearance.

Insulin tolerance test (ITT) was performed to evaluate the role of FoxO1^MPOA^ in insulin sensitivity. Following a 4-hour fast (starting at 9 am), with *ad libitum* access to water, baseline blood glucose levels were measured from tail vein samples using a glucometer. Mice received an intraperitoneal injection of insulin (0.5 U/kg; #0002-75-1001, Eli Lilly and Company, Indianapolis, IN) to induce hypoglycemia. Blood glucose levels were subsequently measured at 15, 30, 60, and 120 minutes post-injection. The temporal glucose measurements were recorded to analyze the dynamic insulin response and assess insulin sensitivity.

### Adulthood FoxO1 MPOA activation

Adult Rosa26-LSL-FoxO1AAA mice underwent bilateral stereotaxic injections into the MPOA with either AAV-CMV-Cre (FoxO1-CA^MPOA^ group; UNC Vector Core; 4×10¹² GC/mL; 200 nL per side) or AAV-CMV-GFP (control group; UNC Vector Core; 4×10¹² GC/mL; 200 nL per side). After recovery, mice were single-housed and maintained on HFD for 10 weeks, during which body weight and food intake were measured weekly. MPOA targeting efficiency and FoxO1AAA overexpression were verified through GFP fluorescence microscopy.

### Ovariectomy (OVX) and hormone supplementation

To investigate the role of ovarian hormones in sex-specific differences, we performed OVX and 17β-estradiol supplementation experiments using control and FoxO1-KO^MPOA^ mice. Eight-week-old female FoxO1^flox/flox^ mice were randomly assigned to either OVX alone or OVX with 17β-estradiol supplementation (OVX+E). For the OVX+E group, a 17β-estradiol pellet (0.025 mg/p, 60-day release, #SE-121, Innovative Research of America, Sarasota, FL) was subcutaneously implanted in the lateral neck region. Additionally, age-matched male FoxO1^flox/flox^ mice also received 17β-estradiol supplementation. During the same surgical procedure, mice in each experimental group underwent bilateral MPOA injections of either AAV-CMV-GFP (control) or AAV-CMV-Cre-GFP (FoxO1-KO^MPOA^). After a 4-day post-surgical recovery period, mice were individually housed and maintained on HFD. Body weight and food intake were monitored at 4-day intervals. Following 30 days of HFD feeding, metabolic parameters of female OVX and male+E mice were assessed using indirect calorimetry in Promethion system as described above.

### Secondary analysis of scRNA-seq results

Single-cell RNA sequencing data from mouse brain MPOA was obtained from NCBI GEO (accession: GSE113576). Data analysis was performed using Seurat v5. Following data acquisition via Read10X, quality control was implemented by excluding nuclei containing >30,000 reads. The filtered dataset underwent log-normalization, and variable features were identified using variance-stabilizing transformation (n=2000 features). Data scaling was performed to standardize gene expression. To mitigate batch effects and technical artifacts, we calculated the percentage of mitochondrial (mt-), ribosomal (Rpl-, Rps-), and immediate-early genes (Fos, Egr1, Npas4). Genes exhibiting strong correlations (|r| > 0.2) with these factors were excluded from variable features. Furthermore, genes expressed in <20 cells were eliminated from analysis.

Dimensionality reduction was achieved through Principal Component Analysis (PCA). Following computation of 50 principal components, the first 15 PCs were selected based on variance contribution visualized via elbow plot. These 15 PCs were utilized to construct a K-nearest neighbors (KNN) graph, followed by cluster identification using the Louvain algorithm. For visualization, two non-linear dimensionality reduction methods were employed: Uniform Manifold Approximation and Projection (UMAP) and t-distributed Stochastic Neighbor Embedding (t-SNE). The t-SNE analysis was performed with perplexity=30 to optimize local-global structure preservation. Cluster-specific marker genes were identified using Seurat, and their expression patterns were visualized through heatmaps, violin plots, and feature plots.

Cell type annotation was performed using two reference-based methods: SingleR with MouseRNAseqData reference dataset and Azimuth with mouse cortex reference. Predictions were validated using average cluster scores, followed by manual annotation refinement. FoxO1 expression analysis was conducted across clusters, and differential expression analysis was performed between FoxO1-positive and FoxO1-negative cells within relevant clusters. Significantly differentially expressed genes associated with FoxO1 expression were identified and filtered for downstream analysis.

### Statistics

Statistical analyses were conducted using GraphPad Prism. The selection of specific statistical methods was determined by the experimental design and is described in the corresponding figure legends. The data is presented as the mean ± standard error of the mean (SEM). A significance threshold of P≤0.05 was employed to define statistical significance.

### Study approval

Care of all animals and procedures were approved by The University of Illinois Chicago Animal Care and Use Committee.

## Author Contributions

P.L. is the main contributor to the study’s conduct, data collection and analysis, data interpretation, and manuscript writing. X.Y., Q.X., L.L.H., V.C.T.I., L.C.S., M.D.M., W.D., L.I., S.S., N.P., N.A., D.D., M.K., C.F., and E.C. contributed to the conduct of the study. C.W.L., G.P., Y.J., C.L., G.S., H.Y., Y.H., T.U., and P.X. contributed to the study design, data interpretation, and manuscript writing.

## Acknowledgements

The authors gratefully acknowledge the Biologic Resources Laboratory at the University of Illinois Chicago for their invaluable assistance with mouse colony maintenance, and the Metabolic Phenotyping Core at the University of Illinois Chicago for performing indirect calorimetry measurements. We thank Dr. Domenico Accili for generously providing the FoxO1-GFP mice and the Vector Core at the University of North Carolina at Chapel Hill for providing AAV vectors.

---

**Figure S1.**
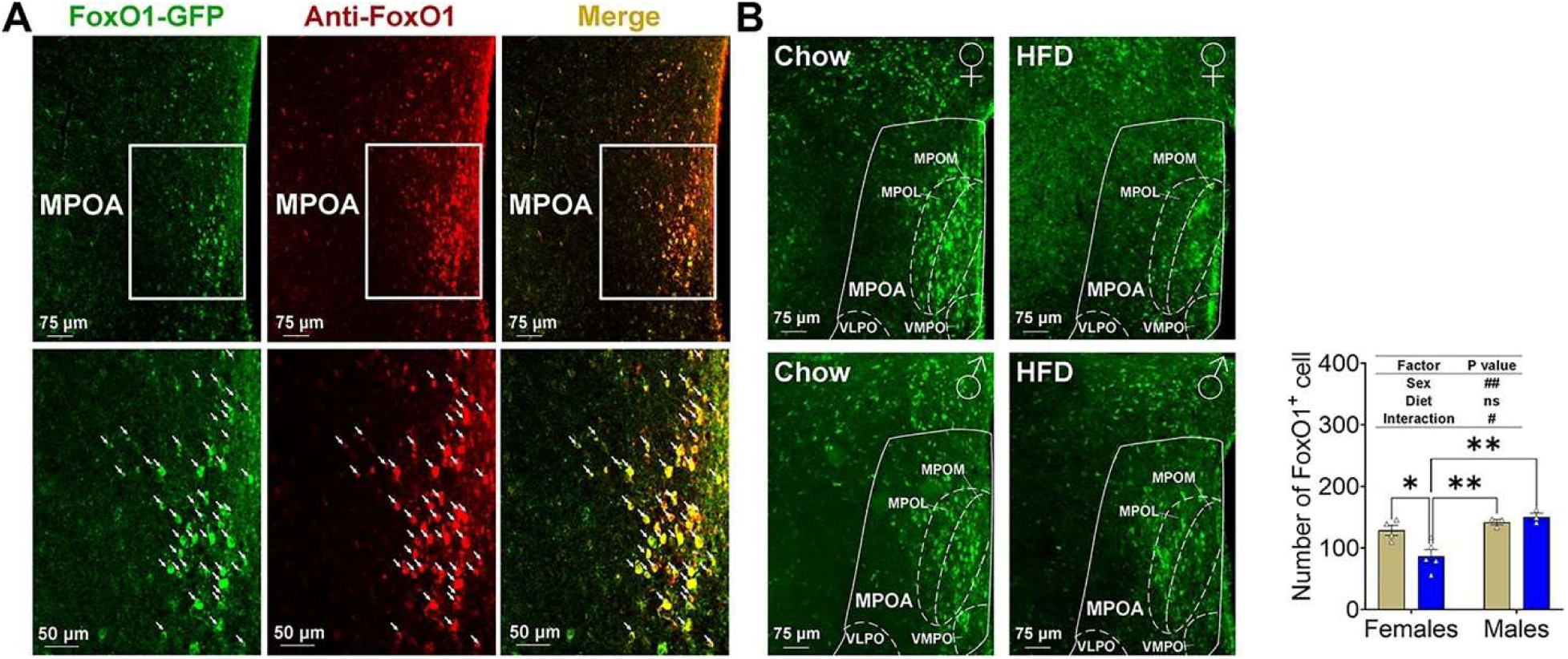
Sex-specific FoxO1 response to diet in the MPOA detected by FoxO1 immunostaining. ***(A)*** Representative images showing the colocalization of FoxO1-GFP and FoxO1 immunofluorescence signals using anti-FoxO1 antibody. White arrowheads point to neurons expressing both GFP and FoxO1. ***(B)*** FoxO1 immunofluorescence and quantification in the MPOA of female or male C57BL/6J mice (8 weeks) after two weeks of either chow diet or HFD feeding are shown. Results are expressed as means ± SEM. (B), #P < 0.05, ##P < 0.01 in the two-way ANOVA analysis, *P < 0.05, **P < 0.01 in the subsequent post hoc Sidak tests.

**Figure S2.**
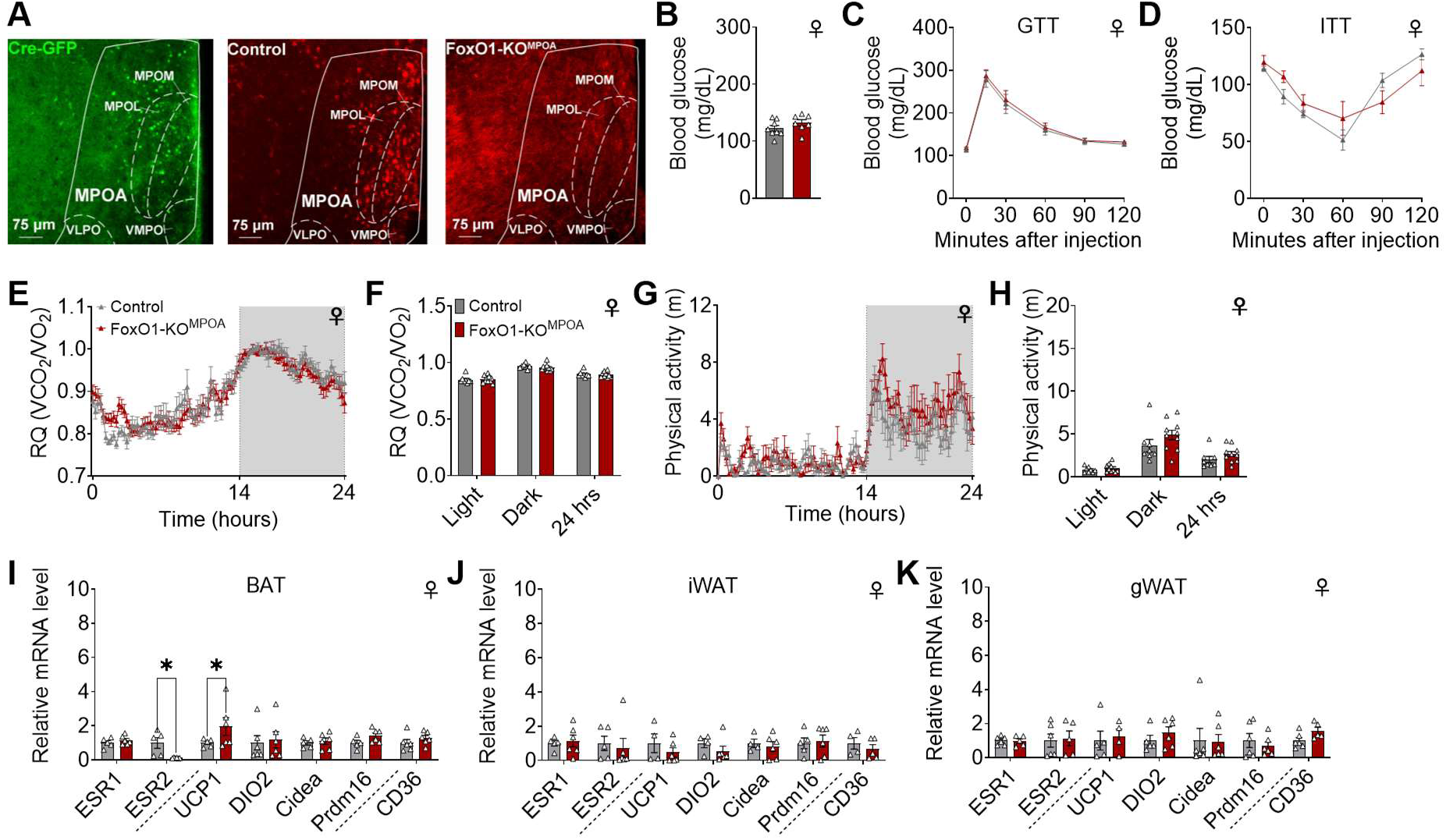
Effect of FoxO1-KO^MPOA^ on female metabolic parameters. ***(A)*** GFP immunoreactivity (left), FoxO1 immunoreactivity in control (middle), and FoxO1-KO^MPOA^ mice. AAV-CMV-Cre-GFP was injected into the MPOA of FoxO1^flox/flox^ mice to generate FoxO1-KO^MPOA^ mice. Control mice were FoxO1^flox/flox^ mice receiving AAV-CMV-GFP virus injections. ***(B)*** Fed blood glucose levels measured 25 weeks after virus injection (n = 8/7). ***(C-D)*** Glucose tolerance tests (C) and insulin tolerance tests (D) were performed 7 and 8 weeks after the virus injection (n = 8/8), using a separate cohort from the body weight recording group. ***(E-H)*** RQ (E), light/dark/24-hour average RQ (F), physical activity (G), and light/dark/24-hour average physical activity (H) in female mice (n = 7/8). ***(I-K)*** mRNA levels of genes in BAT (I), iWAT (J), and gWAT (K) of female mice 25 weeks after virus injections. Measured genes include estrogen receptors (ESR1 and ESR2), thermogenic genes (UCP1, Dio2, Cidea, PRDM16), and a fatty acid sensor and transporter (CD36, n = 6/6). Results are shown as means ± SEM. (I) \**P* < 0.05 in unpaired *t* tests.

**Figure S3.**
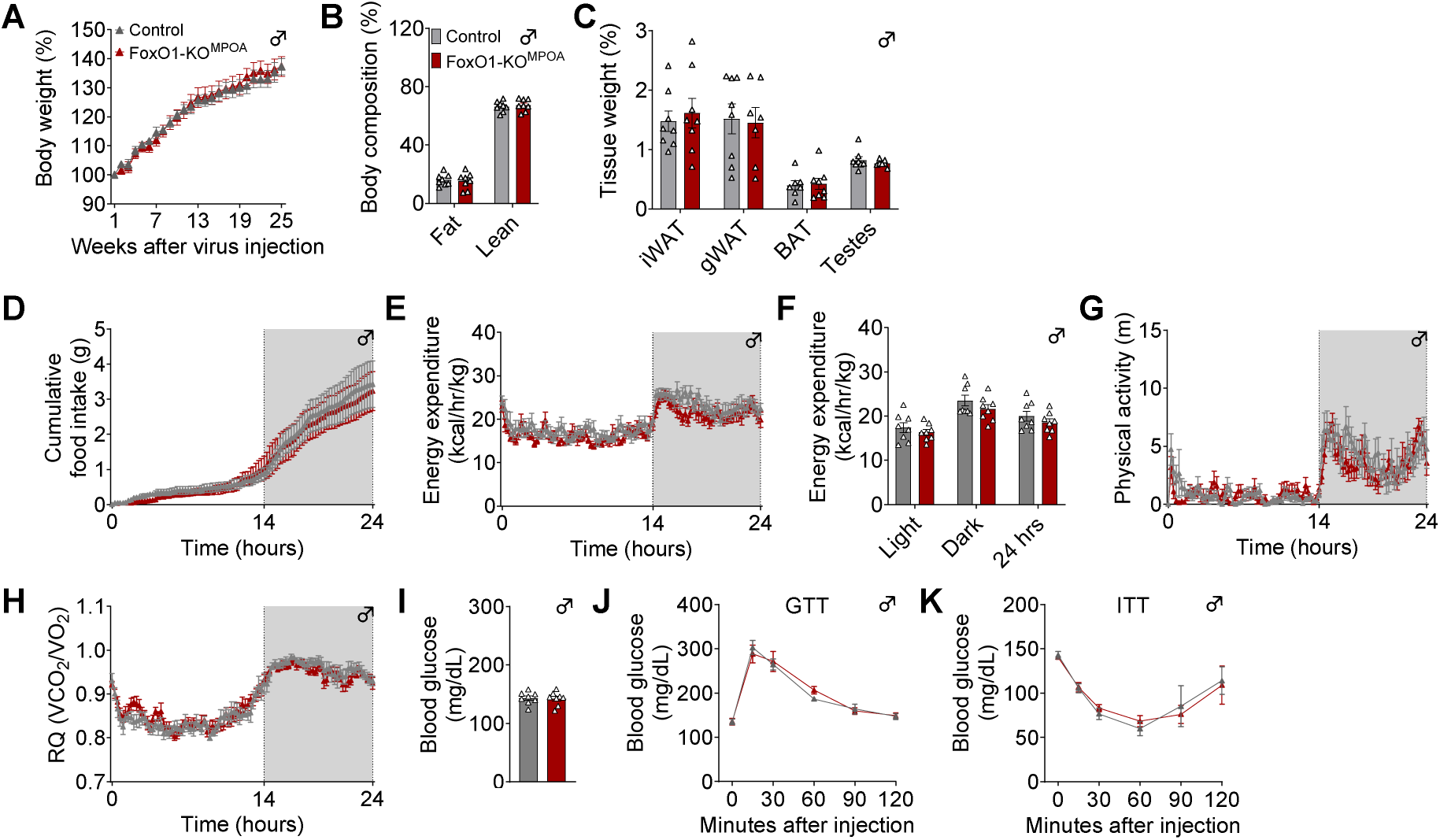
Glucose and energy homeostasis unaffected by FoxO1-KO^MPOA^ in male mice. ***(A)*** Body weight percentage compared to baseline one week after surgery. Male control and FoxO1-KO^MPOA^ mice received virus injections at 8 weeks of age and were fed a chow diet for 25 weeks (n = 8/8). ***(B-C)*** Body composition as percentage of total body weight (B) and tissue index as percentage of body weight (C) in male mice 25 weeks after virus injection (n = 8/8). ***(D-H)*** Cumulative food intake (D), energy expenditure (E), average energy expenditure during light/dark cycles and over 24 hours (F), physical activity (G), and RQ (H) in male mice (n = 8/8). ***(I)*** Fed blood glucose levels measured 25 weeks after virus injection (n = 8/8). ***(J-K)*** Glucose tolerance tests (J) and insulin tolerance tests (K) performed at 7 and 8 weeks after virus injection (n = 8/8). These tests were conducted in a separate cohort from the body weight measurements. Results are presented as means ± SEM.

**Figure S4.**
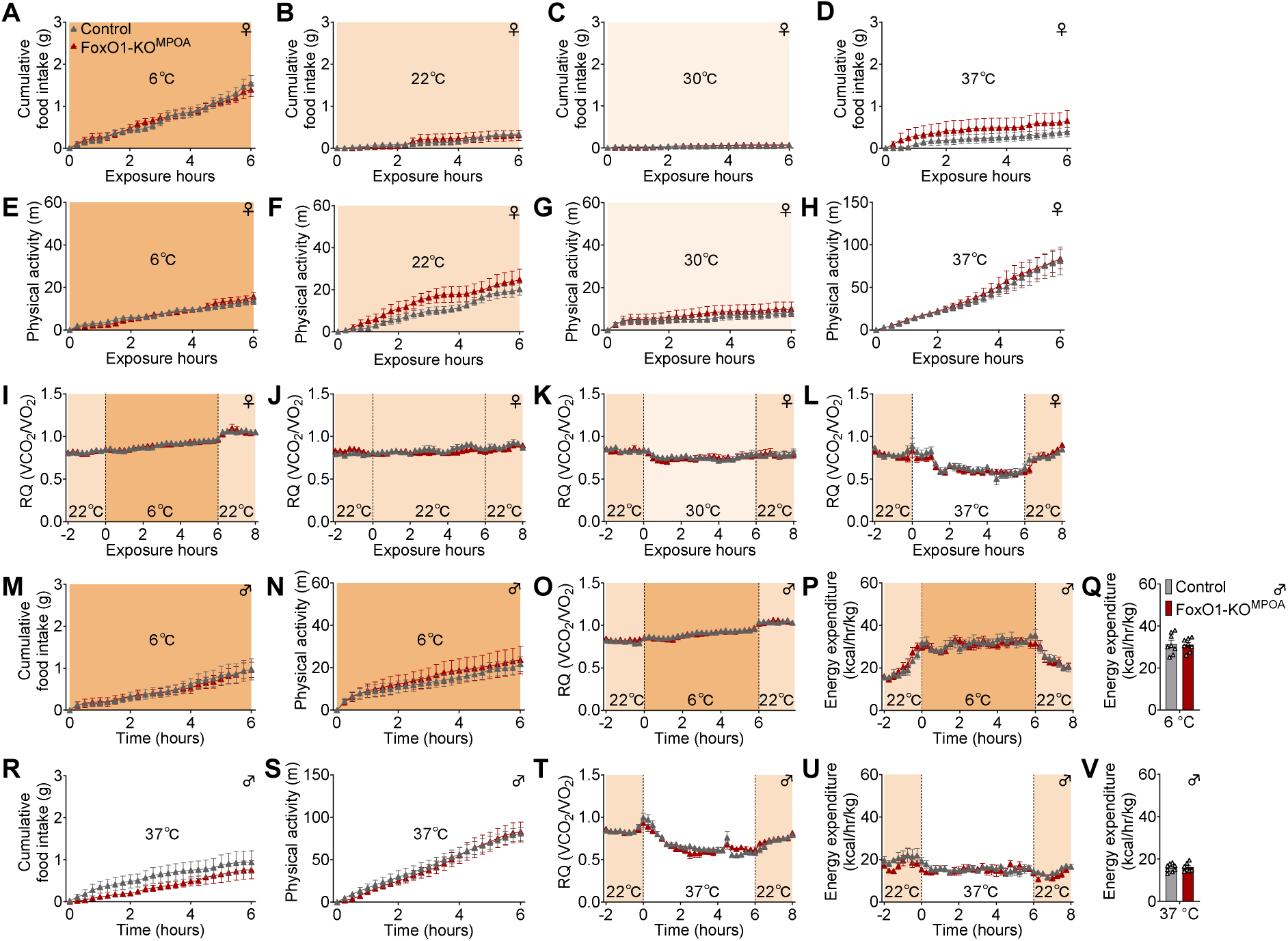
Effects of FoxO1-KO^MPOA^ on metabolic parameters during acute temperature challenges in male and female mice. ***(A–D)*** Cumulative food intake of female mice during 6-hour temperature exposures at 6°C (A, n = 9/8), 22°C (B, n = 5/7), 30°C (C, n = 9/10), and 37°C (D, n = 9/10). ***(E–H)*** Physical activity of female mice during 6-hour temperature exposures at 6°C (E, n = 9/8), 22°C (F, n = 5/7), 30°C (G, n = 9/10), and 37°C (H, n = 9/10). ***(I–L)*** RQ of female mice during 6-hour temperature exposures at 6°C (I, n = 9/8), 22°C (J, n = 5/7), 30°C (K, n = 9/10), and 37°C (L, n = 9/10). ***(M–Q)*** Cumulative food intake (M), physical activity (N), RQ (O), energy expenditure (P), and average energy expenditure (Q) of male mice during 6-hour temperature exposure at 6°C (n = 8/9). ***(R–V)*** Cumulative food intake (R), physical activity (S), RQ (T), energy expenditure (U), and average energy expenditure (V) of male mice during 6-hour temperature exposure at 37°C (n = 9/10). Results are displayed as means ± SEM.

**Figure S5.**
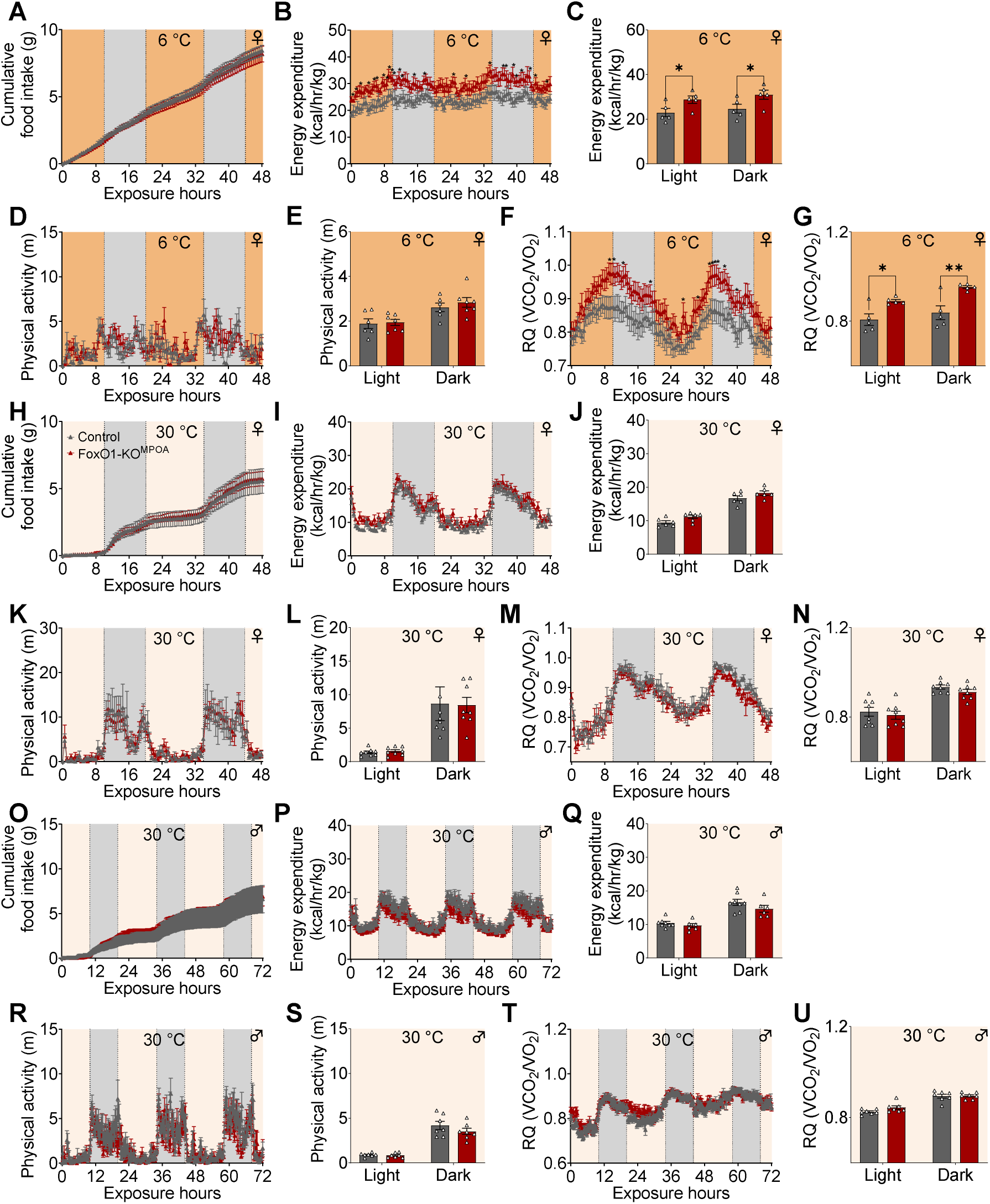
Effects of FoxO1-KO^MPOA^ on energy metabolism during chronic thermoneutral and cold Exposure. ***(A-G)*** Female mice during 48-hour cold exposure were analyzed for cumulative food intake (A, n = 7/8), energy expenditure (B, n = 5/5), light/dark average energy expenditure (C, n = 5/5), 30-minute physical activity (D, n = 6/7), light/dark average physical activity (E, n = 6/7), RQ (F, n = 5/5), and light/dark average RQ (G, n = 5/5). ***(H-N)*** Female mice during 48-hour thermoneutral exposure (n = 7/8) were measured for cumulative food intake (H), energy expenditure (I), light/dark average energy expenditure (J), 30-minute physical activity (K), light/dark average physical activity (L), RQ (M), and light/dark average RQ (N). ***(O-U)*** Male mice during 72-hour thermoneutral exposure (n = 7/7) were assessed for cumulative food intake (O), energy expenditure (P), light/dark average energy expenditure (Q), 30-minute physical activity (R), light/dark average physical activity (S), RQ (T), and light/dark average RQ (U). All measurements were conducted 8 weeks post-virus injection in mice with comparable body weight and lean mass. Data are presented as means ± SEM. Statistical significance was determined by two-way ANOVA with post hoc Sidak tests for panels B-C and F-G (\**P* < 0.05, \*\**P* < 0.01).

**Figure S6.**
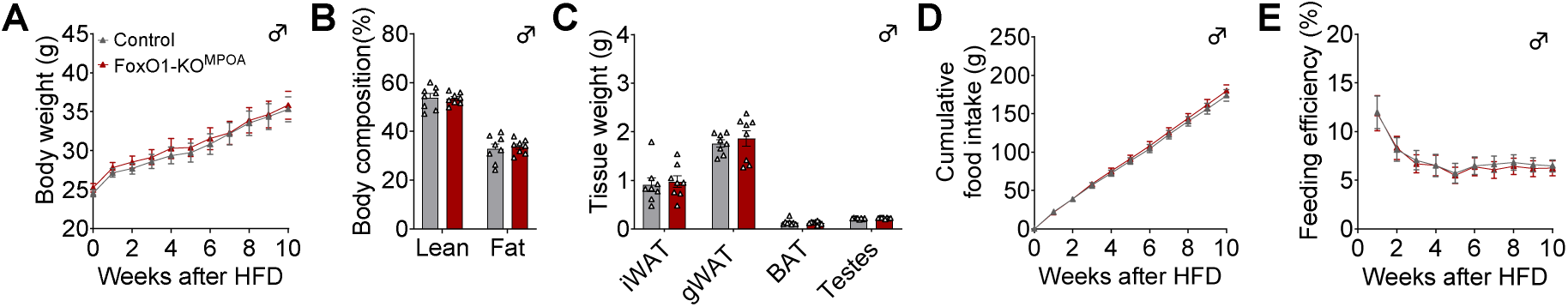
Energy homeostasis unchanged in male FoxO1-KO^MPOA^ mice on HFD. ***(A-C)*** Male mice were fed HFD for 10 weeks (n = 8 per group). Measurements show body weight (A), body composition as percentage of total weight (B), and tissue weights (C). ***(D-E)*** Food intake and feeding efficiency in male control and FoxO1-KO^MPOA^ mice (n = 10 per group). Cumulative food intake (D) and feeding efficiency (E) were monitored during 10 weeks of HFD feeding, initiated 2 weeks post-virus injection at 10 weeks of age. Feeding efficiency represents the ratio of body weight change to cumulative food intake. Data are presented as means ± SEM.

**Figure S7.**
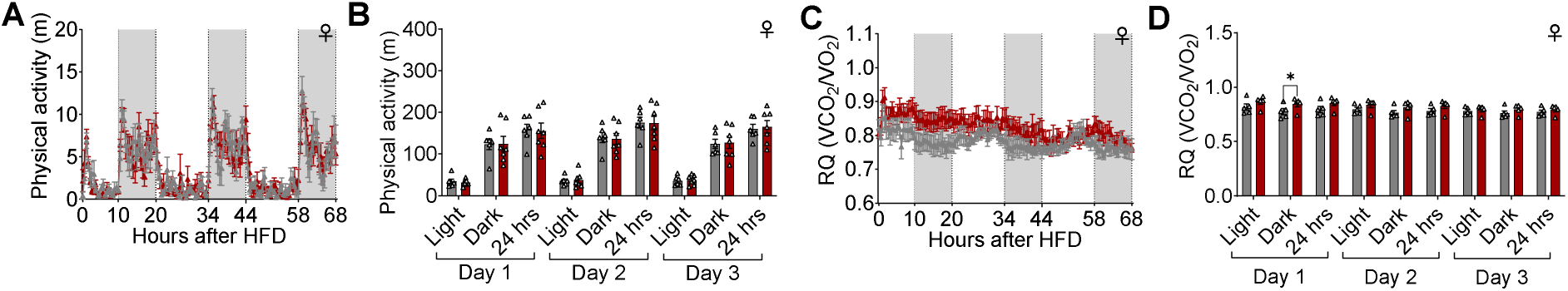
Physical activity and RQ remain unchanged in female FoxO1-KO^MPOA^ mice on HFD. ***(A-B)*** Physical activity measurements were recorded (A) and averaged across light/dark cycles and 24-hour periods (B, n = 6-7 mice per group). ***(C-D)*** Respiratory quotient (RQ) was measured continuously (C) and averaged across light/dark cycles and 24-hour periods (D). Female control and FoxO1-KO^MPOA^ mice were acclimated to metabolic chambers for 2 days on chow diet before transitioning to HFD (n = 5 mice per group). Data are presented as mean ± SEM. (D) \**P* < 0.05, two-way ANOVA with Sidak’s post hoc test.

**Figure S8.**
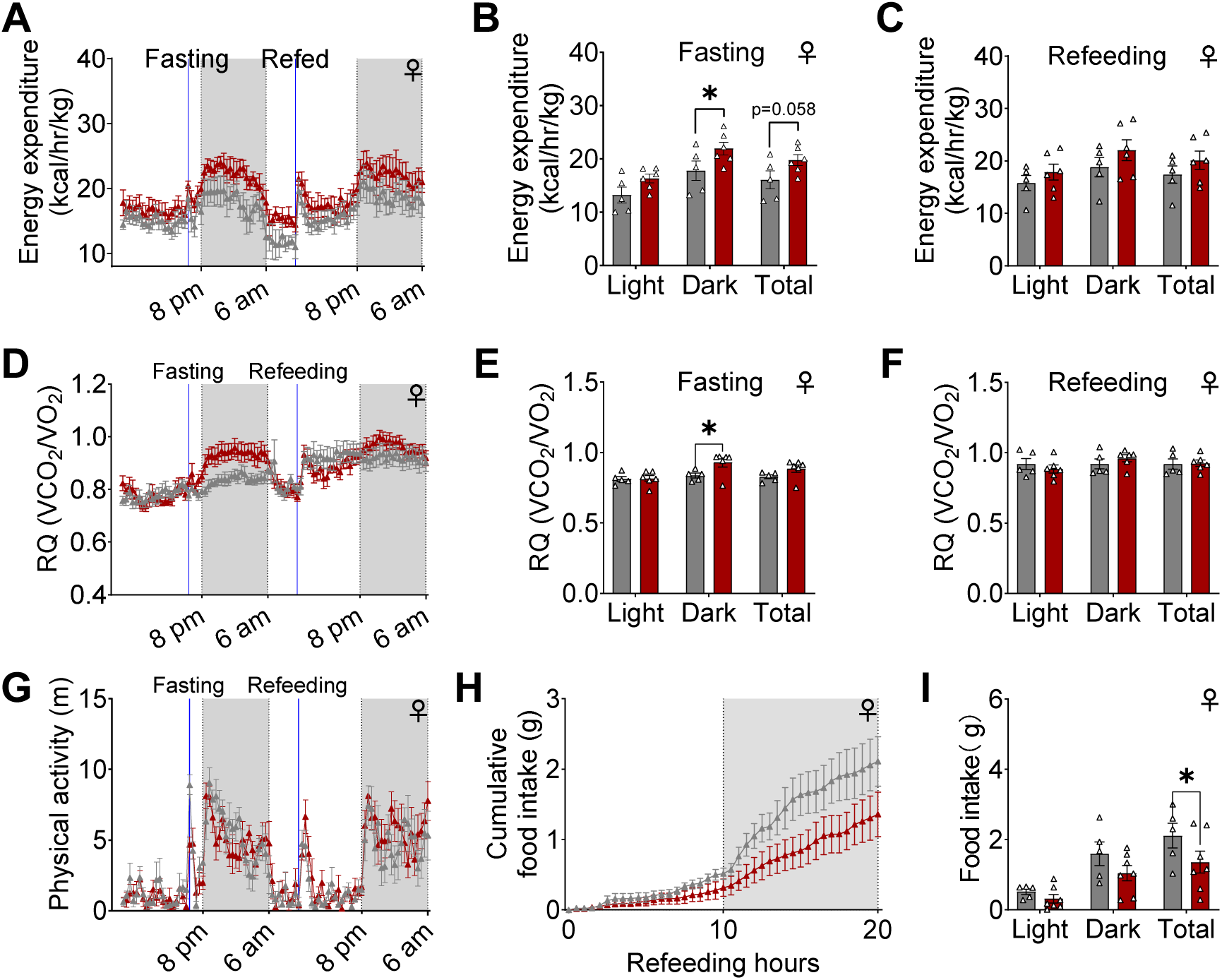
FoxO1-KO^MPOA^ increases energy expenditure during fasting and reduces fast-induced refeeding in female mice. ***(A-I)*** Female control and FoxO1-KO^MPOA^ mice were acclimated to Sable Promethion System for 2 days with chow diet access, followed by overnight fasting and subsequent refeeding with chow diet (n=5-6 per group). (A) Time course of energy expenditure. (B,C) Average energy expenditure during (B) fasting and (C) refeeding periods. (D-F) RQ measurements (D) over time with averages during (E) fasting and (F) refeeding. (G) Physical activity levels. (H,I) Food intake shown as (H) cumulative intake and (I) total intake during refeeding period. Data shown as mean ± SEM. (B, E, I) *P < 0.05, two-way ANOVA with Sidak’s post-hoc test.

**Figure S9.**
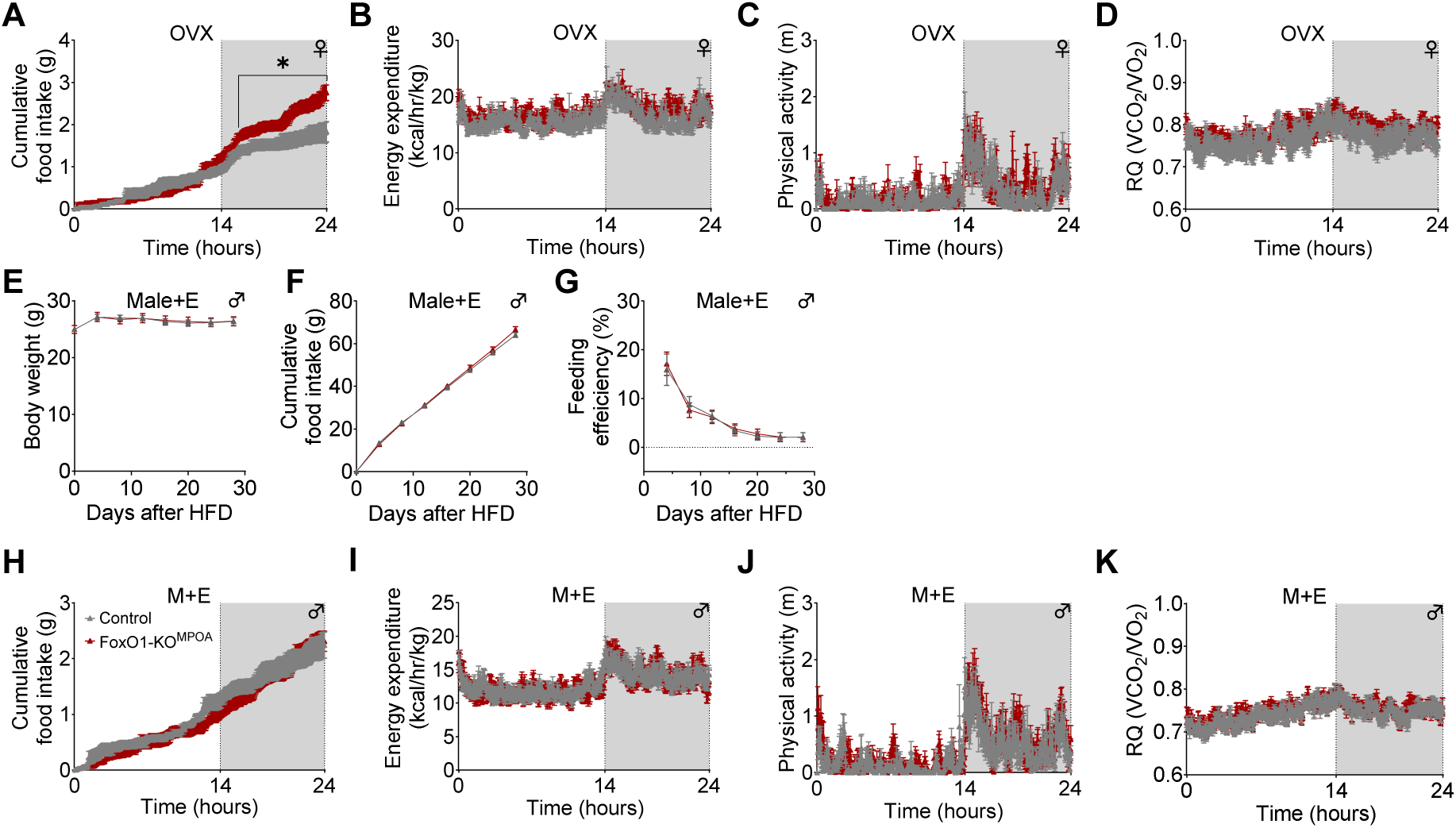
Effects of ovariectomy and 17β-estradiol supplementation on the FoxO1-KO^MPOA^’s impact on energy homeostasis in mice. *(**A-D**)* Analysis of ovariectomized female control and FoxO1-KO^MPOA^ mice (n = 6-8) measuring cumulative food intake (A), energy expenditure (B), physical activity (C), and RQ (D). *(**E-G**)* Male mice were implanted subcutaneously with 17β-estradiol pellets (Male+E, 0.025 mg/pellet, 60-day release, n = 8-9) and monitored for body weight (E), cumulative food intake (F), and feeding efficiency (G). HFD feeding began 4 days post-virus injection and E pellet implantation at 8 weeks of age. *(**H-K**)* Analysis of E pellet-implanted male mice (Male+E, n = 8-9) measuring cumulative food intake (H), energy expenditure (I), physical activity (J), and RQ (K). Mice were fed HFD starting 4 days after virus injection with concurrent OVX or E pellet implantation at 8 weeks of age. Metabolic measurements were conducted using the Sable Promethion System after 30 days on HFD. Data represent means ± SEM. (D) *P < 0.05 by two-way ANOVA with post hoc Sidak tests.

**Figure S10.**
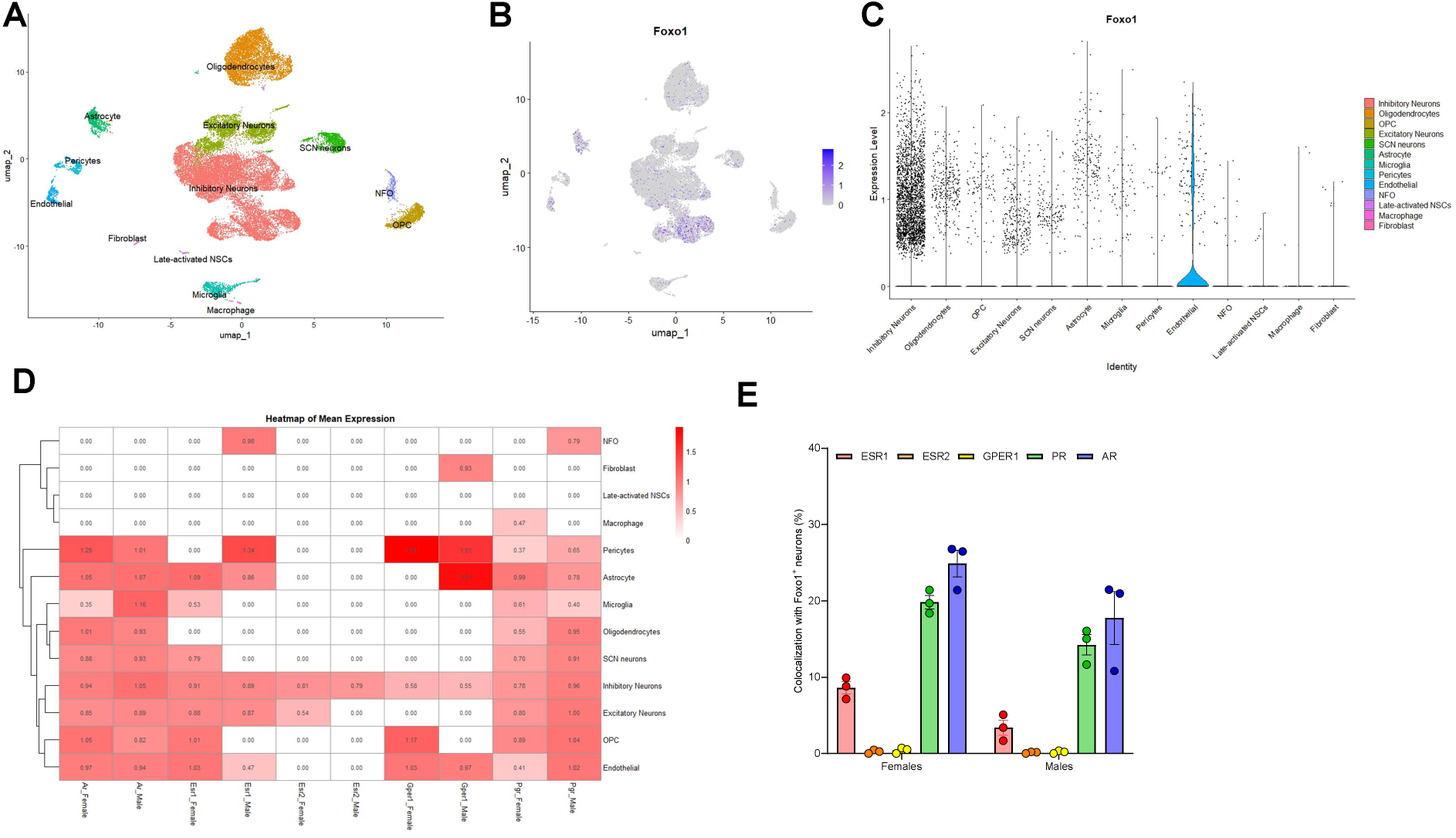
Cell type-specific expression and hormone receptor colocalization of FoxO1 in the MPOA. ***(A)*** t-distributed stochastic neighbor embedding (tSNE) visualization of MPOA cell clusters identified and annotated using SingleR and Azimuth algorithms. OPC: oligodendrocyte progenitor cells; SCN: Suprachiasmatic nucleus neurons; NFO: newly formed oligodendrocytes; NSCs: neural stem cells. ***(B-C)*** Distribution of FoxO1 expression across identified cell populations shown as feature plot (B) and violin plot (C). ***(D)*** Sex hormone receptor expression in FoxO1+ Cells. ESR1, estrogen receptor 1; ESR2, estrogen receptor 2; GPER1, G protein-coupled estrogen receptor 1; PR, progesterone receptor; AR, androgen receptor. ***(E)*** Percentage of FoxO1+ inhibitory neurons expressing individual sex hormone receptor genes. Data represent means ± SEM.

## Reference

1 Chang, Y. & Durante, K. M. Why consumers have everything but happiness: An evolutionary mismatch perspective. Curr Opin Psychol 46, 101347, doi:10.1016/j.copsyc.2022.101347 (2022).

2 Gluckman, P. D., Beedle, A., Buklijas, T., Low, F. & Hanson, M. A. (Oxford University Press, Oxford, 2016).

3 Berenbaum, F., Wallace, I. J., Lieberman, D. E. & Felson, D. T. Modern-day environmental factors in the pathogenesis of osteoarthritis. Nat Rev Rheumatol 14, 674–681, doi:10.1038/s41584-018-0073-x (2018).

4 Ocobock, C. & Niclou, A. Commentary-fat but fit…and cold? Potential evolutionary and environmental drivers of metabolically healthy obesity. Evol Med Public Health 10, 400–408, doi:10.1093/emph/eoac030 (2022).

5 An, R., Ji, M. & Zhang, S. Global warming and obesity: a systematic review. Obes Rev 19, 150–163, doi:10.1111/obr.12624 (2018).

6 Kanazawa, S. Does global warming contribute to the obesity epidemic? Environ Res 182, 108962, doi:10.1016/j.envres.2019.108962 (2020).

7 Eijkelenboom, A. & Burgering, B. M. FOXOs: signalling integrators for homeostasis maintenance. Nat Rev Mol Cell Biol 14, 83–97, doi:10.1038/nrm3507 (2013).

8 Ogg, S. et al. The Fork head transcription factor DAF-16 transduces insulin-like metabolic and longevity signals in C. elegans. Nature 389, 994–999, doi:10.1038/40194 (1997).

9 Lin, K., Dorman, J. B., Rodan, A. & Kenyon, C. daf-16: An HNF-3/forkhead family member that can function to double the life-span of Caenorhabditis elegans. Science 278, 1319–1322, doi:10.1126/science.278.5341.1319 (1997).

10 Kenyon, C., Chang, J., Gensch, E., Rudner, A. & Tabtiang, R. A C. elegans mutant that lives twice as long as wild type. Nature 366, 461–464, doi:10.1038/366461a0 (1993).

11 Gross, D. N., van den Heuvel, A. P. & Birnbaum, M. J. The role of FoxO in the regulation of metabolism. Oncogene 27, 2320–2336, doi:10.1038/onc.2008.25 (2008).

12 Postic, C., Dentin, R. & Girard, J. Role of the liver in the control of carbohydrate and lipid homeostasis. Diabetes Metab 30, 398–408, doi:10.1016/s1262-3636(07)70133-7 (2004).

13 Bastie, C. C. et al. FoxO1 stimulates fatty acid uptake and oxidation in muscle cells through CD36-dependent and -independent mechanisms. J Biol Chem 280, 14222–14229, doi:10.1074/jbc.M413625200 (2005).

14 Kamei, Y. et al. A forkhead transcription factor FKHR up-regulates lipoprotein lipase expression in skeletal muscle. FEBS Lett 536, 232–236, doi:10.1016/s0014-5793(03)00062-0 (2003).

15 Zhang, X. et al. Cold-induced FOXO1 nuclear transport aids cold survival and tissue storage. Nat Commun 15, 2859, doi:10.1038/s41467-024-47095-w (2024).

16 Plum, L. et al. The obesity susceptibility gene Cpe links FoxO1 signaling in hypothalamic pro-opiomelanocortin neurons with regulation of food intake. Nat Med 15, 1195–1201, doi:10.1038/nm.2026 (2009).

17 Kim, M. S. et al. Role of hypothalamic Foxo1 in the regulation of food intake and energy homeostasis. Nat Neurosci 9, 901–906, doi:10.1038/nn1731 (2006).

18 Ren, H. et al. FoxO1 target Gpr17 activates AgRP neurons to regulate food intake. Cell 149, 1314–1326, doi:10.1016/j.cell.2012.04.032 (2012).

19 Kitamura, T. et al. Forkhead protein FoxO1 mediates Agrp-dependent effects of leptin on food intake. Nat Med 12, 534–540, doi:10.1038/nm1392 (2006).

20 Kim, K. W. et al. FOXO1 in the ventromedial hypothalamus regulates energy balance. J Clin Invest 122, 2578–2589, doi:10.1172/JCI62848 (2012).

21 Doan, K. V. et al. FoxO1 in dopaminergic neurons regulates energy homeostasis and targets tyrosine hydroxylase. Nat Commun 7, 12733, doi:10.1038/ncomms12733 (2016).

22 Uchida, Y., Onishi, K., Tokizawa, K. & Nagashima, K. Regional differences of cFos immunoreactive cells in the preoptic areas in hypothalamus associated with heat and cold responses in mice. Neurosci Lett 665, 130–134, doi:10.1016/j.neulet.2017.11.053 (2018).

23 Tansey, E. A. & Johnson, C. D. Recent advances in thermoregulation. Adv Physiol Educ 39, 139–148, doi:10.1152/advan.00126.2014 (2015).

24 Yu, S., Francois, M., Huesing, C. & Munzberg, H. The Hypothalamic Preoptic Area and Body Weight Control. Neuroendocrinology 106, 187–194, doi:10.1159/000479875 (2018).

25 Hamilton, C. L. & Brobeck, J. R. Food Intake and Temperature Regulation in Rats with Rostral Hypothalamic Lesions. Am J Physiol 207, 291–297, doi:10.1152/ajplegacy.1964.207.2.291 (1964).

26 Andersson, B. & Larsson, B. Influence of local temperature changes in the preoptic area and rostral hypothalamus on the regulation of food and water intake. Acta Physiol Scand 52, 75–89, doi:10.1111/j.1748-1716.1961.tb02203.x (1961).

27 Hamilton, C. L. Interactions of food intake and temperature regulation in the rat. J Comp Physiol Psychol 56, 476–488, doi:10.1037/h0046241 (1963).

28 Rodriguez-Cuenca, S. et al. Sex-dependent thermogenesis, differences in mitochondrial morphology and function, and adrenergic response in brown adipose tissue. J Biol Chem 277, 42958–42963, doi:10.1074/jbc.M207229200 (2002).

29 Hrvatin, S. et al. Neurons that regulate mouse torpor. Nature, doi:10.1038/s41586-020-2387-5 (2020).

30 Takahashi, T. M. et al. A discrete neuronal circuit induces a hibernation-like state in rodents. Nature, doi:10.1038/s41586-020-2163-6 (2020).

31 Zhang, Z. et al. Estrogen-sensitive medial preoptic area neurons coordinate torpor in mice. Nat Commun 11, 6378, doi:10.1038/s41467-020-20050-1 (2020).

32 Kuo, T. et al. Identification of C2CD4A as a human diabetes susceptibility gene with a role in beta cell insulin secretion. Proc Natl Acad Sci U S A 116, 20033–20042, doi:10.1073/pnas.1904311116 (2019).

33 da Conceicao, E. P. S., Morrison, S. F., Cano, G., Chiavetta, P. & Tupone, D. Median preoptic area neurons are required for the cooling and febrile activations of brown adipose tissue thermogenesis in rat. Sci Rep 10, 18072, doi:10.1038/s41598-020-74272-w (2020).

34 Hankenson, F. C., Marx, J. O., Gordon, C. J. & David, J. M. Effects of Rodent Thermoregulation on Animal Models in the Research Environment. Comp Med 68, 425–438, doi:10.30802/AALAS-CM-18-000049 (2018).

35 Jash, S., Banerjee, S., Lee, M. J., Farmer, S. R. & Puri, V. CIDEA Transcriptionally Regulates UCP1 for Britening and Thermogenesis in Human Fat Cells. iScience 20, 73–89, doi:10.1016/j.isci.2019.09.011 (2019).

36 Kajimura, S. et al. Initiation of myoblast to brown fat switch by a PRDM16-C/EBP-beta transcriptional complex. Nature 460, 1154–1158, doi:10.1038/nature08262 (2009).

37 Putri, M. et al. CD36 is indispensable for thermogenesis under conditions of fasting and cold stress. Biochem Biophys Res Commun 457, 520–525, doi:10.1016/j.bbrc.2014.12.124 (2015).

38 Xu, Y. et al. Distinct hypothalamic neurons mediate estrogenic effects on energy homeostasis and reproduction. Cell Metab 14, 453–465, doi:10.1016/j.cmet.2011.08.009 (2011).

39 Ouyang, W. et al. Novel Foxo1-dependent transcriptional programs control T(reg) cell function. Nature 491, 554–559, doi:10.1038/nature11581 (2012).

40 Moffitt, J. R., et al. Molecular, spatial, and functional single-cell profiling of the hypothalamic preoptic region. Science 362, doi:10.1126/science.aau5324 (2018).

41 Heinrich, G., Meece, K., Wardlaw, S. L. & Accili, D. Preserved energy balance in mice lacking FoxO1 in neurons of Nkx2.1 lineage reveals functional heterogeneity of FoxO1 signaling within the hypothalamus. Diabetes 63, 1572–1582, doi:10.2337/db13-0651 (2014).

42 Tsuneoka, Y. Molecular neuroanatomy of the mouse medial preoptic area with reference to parental behavior. Anat Sci Int 94, 39–52, doi:10.1007/s12565-018-0468-4 (2019).

43 Qian, S. et al. A temperature-regulated circuit for feeding behavior. Nat Commun 13, 4229, doi:10.1038/s41467-022-31917-w (2022).

44 Yang, S. et al. An mPOA-ARC(AgRP) pathway modulates cold-evoked eating behavior. Cell Rep 36, 109502, doi:10.1016/j.celrep.2021.109502 (2021).

45 Herz, C. T. et al. Sex differences in brown adipose tissue activity and cold-induced thermogenesis. Mol Cell Endocrinol 534, 111365, doi:10.1016/j.mce.2021.111365 (2021).

46 Keuper, M. & Jastroch, M. The good and the BAT of metabolic sex differences in thermogenic human adipose tissue. Mol Cell Endocrinol 533, 111337, doi:10.1016/j.mce.2021.111337 (2021).

47 Harshaw, C., Culligan, J. J. & Alberts, J. R. Sex differences in thermogenesis structure behavior and contact within huddles of infant mice. PLoS One 9, e87405, doi:10.1371/journal.pone.0087405 (2014).

48 Gomez-Garcia, I., Trepiana, J., Fernandez-Quintela, A., Giralt, M. & Portillo, M. P. Sexual Dimorphism in Brown Adipose Tissue Activation and White Adipose Tissue Browning. Int J Mol Sci 23, doi:10.3390/ijms23158250 (2022).

49 Fernandez-Pena, C., Reimundez, A., Viana, F., Arce, V. M. & Senaris, R. Sex differences in thermoregulation in mammals: Implications for energy homeostasis. Front Endocrinol (Lausanne) 14, 1093376, doi:10.3389/fendo.2023.1093376 (2023).

50 Schuur, E. R. et al. Ligand-dependent interaction of estrogen receptor-alpha with members of the forkhead transcription factor family. J Biol Chem 276, 33554–33560, doi:10.1074/jbc.M105555200 (2001).

51 Yan, H. et al. Estrogen Protects Cardiac Function and Energy Metabolism in Dilated Cardiomyopathy Induced by Loss of Cardiac IRS1 and IRS2. Circ Heart Fail 15, e008758, doi:10.1161/CIRCHEARTFAILURE.121.008758 (2022).

52 Yan, H. et al. Estrogen Improves Insulin Sensitivity and Suppresses Gluconeogenesis via the Transcription Factor Foxo1. Diabetes 68, 291–304, doi:10.2337/db18-0638 (2019).

53 Lengyel, F. et al. Effect of estrogen and inhibition of phosphatidylinositol-3 kinase on Akt and FOXO1 in rat uterus. Steroids 72, 422–428, doi:10.1016/j.steroids.2007.03.001 (2007).

54 Fuller-Jackson, J. P., Dordevic, A. L., Clarke, I. J. & Henry, B. A. Effect of sex and sex steroids on brown adipose tissue heat production in humans. Eur J Endocrinol 183, 343–355, doi:10.1530/EJE-20-0184 (2020).

55 Taniguchi, H., Hashimoto, Y., Dowaki, N. & Nirengi, S. Association of brown adipose tissue activity with circulating sex hormones and fibroblast growth factor 21 in the follicular and luteal phases in young women. J Physiol Anthropol 43, 23, doi:10.1186/s40101-024-00371-6 (2024).

56 Hoyenga, K. B. & Hoyenga, K. T. Gender and energy balance: sex differences in adaptations for feast and famine. Physiol Behav 28, 545–563, doi:10.1016/0031-9384(82)90153-6 (1982).

57 Zarulli, V. et al. Women live longer than men even during severe famines and epidemics. Proc Natl Acad Sci U S A 115, E832–E840, doi:10.1073/pnas.1701535115 (2018).

58 Pietrobelli, A. et al. Sexual dimorphism in the energy content of weight change. Int J Obes Relat Metab Disord 26, 1339–1348, doi:10.1038/sj.ijo.0802065 (2002).

59 Wu, B. N. & O’Sullivan, A. J. Sex differences in energy metabolism need to be considered with lifestyle modifications in humans. J Nutr Metab 2011, 391809, doi:10.1155/2011/391809 (2011).

60 Stubbins, R. E., Holcomb, V. B., Hong, J. & Nunez, N. P. Estrogen modulates abdominal adiposity and protects female mice from obesity and impaired glucose tolerance. Eur J Nutr 51, 861–870, doi:10.1007/s00394-011-0266-4 (2012).

61 D’Eon, T. M. et al. Estrogen regulation of adiposity and fuel partitioning. Evidence of genomic and non-genomic regulation of lipogenic and oxidative pathways. J Biol Chem 280, 35983–35991, doi:10.1074/jbc.M507339200 (2005).

62 Omotola, O., Legan, S., Slade, E., Adekunle, A. & Pendergast, J. S. Estradiol regulates daily rhythms underlying diet-induced obesity in female mice. Am J Physiol Endocrinol Metab 317, E1172–E1181, doi:10.1152/ajpendo.00365.2019 (2019).

63 Skinner, D. C., Caraty, A. & Allingham, R. Unmasking the progesterone receptor in the preoptic area and hypothalamus of the ewe: no colocalization with gonadotropin-releasing neurons. Endocrinology 142, 573–579, doi:10.1210/endo.142.2.7956 (2001).

64 Diep, C. H. et al. Progesterone receptors induce FOXO1-dependent senescence in ovarian cancer cells. Cell Cycle 12, 1433–1449, doi:10.4161/cc.24550 (2013).

65 Stelmanska, E. & Sucajtys-Szulc, E. Enhanced food intake by progesterone-treated female rats is related to changes in neuropeptide genes expression in hypothalamus. Endokrynol Pol 65, 46–56, doi:10.5603/EP.2014.0007 (2014).

66 Holmberg, E. et al. Allopregnanolone involvement in feeding regulation, overeating and obesity. Front Neuroendocrinol 48, 70–77, doi:10.1016/j.yfrne.2017.07.002 (2018).

67 Hervey, E. & Hervey, G. R. The effects of progesterone on body weight and composition in the rat. J Endocrinol 37, 361–381, doi:10.1677/joe.0.0370361 (1967).

68 Ashby, J. P., Shirling, D. & Baird, J. D. Differential changes in body composition during growth and progesterone treatment in intact female rats. J Endocrinol 93, 391–395, doi:10.1677/joe.0.0930391 (1982).

69 Bookout, A. L. & Mangelsdorf, D. J. Quantitative real-time PCR protocol for analysis of nuclear receptor signaling pathways. Nucl Recept Signal 1, e012, doi:10.1621/nrs.01012 (2003).

70 Kimball, A. et al. The allopregnanolone to progesterone ratio across the menstrual cycle and in menopause. Psychoneuroendocrinology 112, 104512, doi:10.1016/j.psyneuen.2019.104512 (2020).

71 Kim, B. K. et al. Composite contributions of cerebrospinal fluid GABAergic neurosteroids, neuropeptide Y and interleukin-6 to PTSD symptom severity in men with PTSD. Neurobiol Stress 12, 100220, doi:10.1016/j.ynstr.2020.100220 (2020).

72 Ye, H. et al. An estrogen-sensitive hypothalamus-midbrain neural circuit controls thermogenesis and physical activity. Sci Adv 8, eabk0185, doi:10.1126/sciadv.abk0185 (2022).

73 Bettscheider, M., Murgatroyd, C. & Spengler, D. Simultaneous DNA and RNA isolation from brain punches for epigenetics. BMC Research Notes 4, 314, doi:10.1186/1756-0500-4-314 (2011).

